# Reviving immunogenic cell death upon targeting TACC3 enhances T-DM1 response in HER2-positive breast cancer

**DOI:** 10.1101/2023.09.12.557273

**Authors:** Mustafa Emre Gedik, Ozge Saatci, Nathaniel Oberholtzer, Meral Uner, Ozge Akbulut, Metin Cetin, Mertkaya Aras, Kubra Ibis, Burcu Caliskan, Erden Banoglu, Stefan Wiemann, Aysegul Uner, Sercan Aksoy, Shikhar Mehrotra, Ozgur Sahin

## Abstract

Immunogenic cell death (ICD), an immune-priming form of cell death, has been shown to be induced by several different anti-cancer therapies. Despite being the first and one of the most successful antibody-drug conjugates (ADCs) approved for refractory HER2-positive breast cancer, little is known if response and resistance to trastuzumab emtansine (T-DM1) involves ICD modulation that can be leveraged to enhance T-DM1 response. Here, we report that T-DM1 induces spindle assembly checkpoint (SAC)-dependent ICD in sensitive cells by inducing eIF2α phosphorylation, surface exposure of calreticulin, ATP and HMGB1 release, and secretion of ICD-related cytokines, all of which are lost in resistance. Accordingly, an ICD-related gene signature correlates with clinical response to T-DM1-containing therapy. We found that transforming acidic coiled-coil containing 3 (TACC3) is overexpressed in T-DM1 resistant cells, and that T-DM1 responsive patients have reduced TACC3 protein while the non-responders exhibited increased TACC3 expression during T-DM1 treatment. Notably, genetic or pharmacological inhibition of TACC3 revives T-DM1-induced SAC activation and induction of ICD markers in vitro. Finally, TACC3 inhibition elicits ICD in vivo shown by vaccination assay, and it potentiates T-DM1 by inducing dendritic cell (DC) maturation and enhancing infiltration of cytotoxic T cells in the human HER2-overexpressing MMTV.f.huHER2#5 (Fo5) transgenic model. Together, our results show that ICD is a key mechanism of action of T-DM1 which is lost in resistance, and that targeting TACC3 restores T-DM1-mediated ICD and overcomes resistance.

**Statement of Significance:** Immunogenic cell death (ICD) is a novel mechanism of T-DM1 cytotoxicity that is lost upon T-DM1 resistance. Targeting TACC3 reinstates T-DM1-induced ICD, thus representing an attractive strategy to overcome T-DM1 resistance.

## Introduction

Immunogenic cell death (ICD) is a specific form of cell death that is characterized by the release of antigenic molecules from tumors, e.g., danger-associated molecular patterns (DAMPs), such as cell surface exposure of calreticulin, ATP and HMGB1 release. This, in turn, leads to secretion of pro-inflammatory cytokines that altogether evoke anti-tumor immune responses upon dendritic cell (DC) maturation and cytotoxic T-cell activation^1,2^. Although ICD has been shown to be a key mechanism of cell death for several different chemotherapies^3,4^ or targeted therapy agents^5^, little is known if loss of ICD can trigger resistance to anti-cancer therapies and if there are therapeutic opportunities to revive ICD and restore drug sensitivity.

Antibody-drug conjugates (ADC) are immunoconjugates containing a monoclonal antibody that is bound to a cytotoxic drug, a so-called payload, with a chemical linker^6^. ADCs are highly effective agents and successfully used in treating many different cancers, including breast cancer. Ado-trastuzumab emtansine (T-DM1) is an FDA-approved iconic ADC of a HER2-targeting antibody (trastuzumab) conjugated to a maytansine derivative (DM1) which depolymerizes microtubules. It is the first and one of the most successful ADCs approved for the treatment of aggressive HER2+ breast cancer with refractory disease^7,8^. Despite its initial clinical success, resistance is common. Resistance to ADCs may involve alterations in the internalization or recycling of the targeted tumor antigen, e.g., HER2, in case of T-DM1, changes in payload efficacy, such as enhanced drug efflux or activation of specific survival mechanisms, as demonstrated by us and others^9–12^. However, it is still not fully uncovered (i) if response to ADCs may involve activation of ICD, (ii) if resistance to ADCs may be driven by loss of ICD, and (iii) if there are specific molecular targets that may be inhibited to reinstate ICD and thus achieve a much stronger and more durable response.

Transforming acidic coiled-coil containing 3 (TACC3) is upregulated in solid tumors and hematologic malignancies and is strongly associated with worse prognosis in several different cancers^13–19^. It is localized to centrosomes as well as microtubules to control spindle stability and microtubule nucleation^20^. We previously showed that TACC3 forms distinct functional interactomes regulating different processes in mitosis and interphase to ensure proliferation and survival of cancer cells with centrosome amplification^21^. Despite its roles in tumor growth of highly aggressive cancers, little is known about the role of TACC3 in regulating the immunogenicity of tumors or in driving resistance to anti-cancer therapies.

In this study, we demonstrated, for the first time, that the T-DM1-induced activation of spindle assembly checkpoint (SAC) triggers ICD and contributes to T-DM1 cytotoxicity, while T-DM1 resistance is characterized by lack of ICD induction. Notably, an ICD-related gene signature correlates with clinical response to T-DM1-containing therapy. We further demonstrated that TACC3 is overexpressed in T-DM1 resistant models and the increase in TACC3 protein after T-DM1 therapy is associated with clinical T-DM1 resistance in neo-adjuvant settings. Targeting TACC3 not only restores T-DM1-induced SAC activation and mitotic cell death, but it also activates ICD hallmarks and leads to secretion of pro-inflammatory cytokines in T-DM1 resistant cells. This ultimately results in DC maturation and infiltration of cytotoxic T cells enhancing T-DM1 response in vivo.

## Materials and Methods

### Cell lines, drugs, and culture conditions

Human breast cancer cell lines, SK-BR-3 and BT-474 and murine mammary tumor cell line, EMT6 were obtained from ATCC (Manassas, VA, USA). T-DM1 resistant (T-DM1R) SK-BR-3 and BT-474 cells were generated previously^9^. EMT6.huHER2 cells were generated from the EMT6 cell line by stable overexpression of human HER2. All the cells were cultured in Dulbecco Modified Eagle Medium (Corning, NY, USA) supplemented with 50 U/ml penicillin/streptomycin, 1% non-essential amino acids and 10% fetal bovine serum (Gibco, MT, USA). Cells were routinely tested for mycoplasma contamination using MycoAlert detection kit (Lonza, NJ, USA) and were authenticated by STR sequencing.

### HER2+ human tumor samples

To analyze the association between TACC3 protein expression and response to T-DM1, IHC staining of TACC3 was performed in matched primary tumor samples from 16 female HER2+ breast cancer patients collected before and after therapy. The patients were categorized as responders if they survived longer than 3 years after diagnosis vs. non-responders if they died within 3 years after diagnosis^22,23^, and the change in TACC3 protein expression as fold change was calculated. The patients were diagnosed between 2010 and 2019 at Hacettepe University School of Medicine, Ankara, Turkey. The study was approved by the Non□Interventional Clinical Research Ethics Committee of Hacettepe University (approval no: 2020/02-40). Informed consent was obtained from all patients.

### Transient transfection with siRNAs and overexpression vectors

siRNAs were purchased from Dharmacon (Lafayette, CO, USA), and the sequences are provided in **Supplementary Table S1**. For cell viability, BT-474 T-DM1R (8×10^3^ cells/well) and SK-BR-3 T-DM1R (6×10^3^ cells/well) cells were seeded in 96-well plates with their growth medium without P/S. Twenty-four hours later, cells were transfected with three different siRNAs targeting TACC3 (Dharmacon, CO, USA) at a final concentration of 20 nM (siTACC3#1: D-004155-03, siTACC3#2: D-004155-02, and siTACC3#3: D-004155-04-0005) using Lipofectamine 2000^TM^ (Invitrogen, CA, USA) transfection reagent as described previously^24^. Twenty-four hours after transfections, BT-474 and SK-BR-3 T-DM1R cells were treated with 15 and 0.06 μg/ml of T-DM1, respectively. Cell viability was measured with Sulforhodamine B (SRB) assay 72 hours after T-DM1 treatment. Empty or TACC3 overexpression vector was given 12 hours before T-DM1 treatment with an amount of 100 ng/well. 72 hours following transfection, cell viability was measured with SRB assay. For immunofluorescence experiments, BT-474 T-DM1R cells (3.5×10^5^ cells/well) and SK-BR-3 T-DM1R cells (3×10^5^ cells/well) were seeded into 6- well plates, and next day transfected with siRNAs targeting TACC3 at a final concentration of 40 nM.

### Isolation of bone marrow-derived cells (BMDCs)

To obtain bone marrow-derived cells, 6-8 weeks old BALB/c mice were euthanized, and the femur and tibia were dissected, cleaned, and placed in a 6-well dish containing IMDM culture media. A mortar was used to crush the bones, releasing the bone marrow contents into the culture media. The bone marrow cells were plated in a 6-well plate at a seeding density of 1 million cells/mL (5 mL total per well). The cells were cultured for 7 days with 10 ng/ml GM-CSF and 10 ng/ml IL-4. Dendritic cell phenotype was confirmed by flow cytometry using CD11c positivity.

### DC maturation and T cell activation assays

The DCs were cultured in conditioned media from EMT6.huHER2 cells treated with T-DM1 alone or in combination with BO-264 for a week, and the expression of maturation markers CD80 and CD86 were analyzed by flow cytometry. For the co-culture experiments, DCs were cultured in the conditioned media from EMT6.huHER2 cells treated with T-DM1 alone or in combination with BO-264 for overnight in the presence of gp100 peptide (1 ug/mL). The next day, CD3+ T cells isolated from gp100 T cell receptor bearing Pmel-1 transgenic mice were labeled with CellTrace Violet (Thermo Fisher, MA, USA) and added to the wells containing gp100-loaded DCs at a ratio of 1:10 (DC:T cell). After 5 days in culture, T cells were collected and analyzed for markers of activation and proliferation. Supernatant from the co-culture was collected and sent for multiplex cytokine analysis (Eve Technologies, Alberta, Canada).

### Flow cytometry

To analyze the expression of DC maturation and T cell markers, the DCs and T cells were trypsinized and fixed with 4% PFA. Staining for cell surface markers was performed by incubating cells with conjugated primary antibodies (**Supplementary Table S2**) diluted at 1:200 ratio in FACS buffer (0.1% BSA in PBS) for 30 min at 4°C. Samples were then processed using LSRFortessa and analyzed with FlowJo software (v10.8.1) (Tree Star, OR, USA).

### ATP secretion and HMGB1 release assays

To measure ATP secretion and HMGB1 release, ENLITEN^®^ ATP assay system and LUMIT™ HMGB11 immunoassay (Promega, WI, USA) were utilized, respectively. Cells were seeded in 96 well plate and treated with drugs for 48 hours. Supernatant was transferred into opaque 96 well plates and analyzed according to the manufacturer’s instruction. The luminescence signal was measured using SpectraMax^®^ i3x (Molecular Devices, CA, USA) microplate reader.

### Cytokine array

Human Cytokine Array C5 (RayBiotech, GA, USA) was performed according to the manufacturer’s instructions. Briefly, BT-474 and SK-BR-3 WT cells or SK-BR-3 T-DM1R cells were treated with T-DM1 only or in combination with BO-264, respectively for 48 hours. Then, supernatants were collected and incubated with the antibody-coated membranes overnight. Next day, they were incubated with biotinylated Ab and labeled with streptavidin. The chemiluminescence imaging was performed using iBright Imaging system (Thermo Fisher, MA, USA), and iBright Analysis software (v(5.1.0)) were used for data analysis.

### In vivo studies

Six-to-eight-week-old female BALB/c or FVB mice were housed with a temperature-controlled and 12- hour light/12-hour dark cycle environment. All the in vivo studies were carried out in accordance with the Institutional Animal Care and Use Committee of the University of South Carolina and Medical University of South Carolina. The human HER2-expressing MMTV.f.huHER2#5 (Fo5) transgenic model was obtained from Genentech under a material transfer agreement (OM-217137 and OR-224086B). Tumor pieces of 2×2 mm in size were transplanted near the MFP of female 6-8 weeks old immunocompetent FVB mice. After the mean tumor volume reached 100 mm^3^, mice were randomly allocated to treatment groups. T-DM1 was given once at a dose of 5 mg/kg, by intravenous (i.v.) injection, while BO-264 was given at a dose of 50 mg/kg, daily, via oral gavage. For testing the effects of combination therapy on tumor growth, mice were treated for 2 weeks, and all mice were sacrificed, and tumors were collected for Western blotting. To analyze immune cell infiltration and serum cytokine profiling, a separate cohort of mice were treated for a week, and then sacrificed and serum samples and tumors were collected. Tumors were processed for multiplex IHC staining while serum samples were sent for multiplex cytokine analysis (Eve Technologies, Alberta, Canada). There was no blinding during in vivo experiments. Sample sizes were determined based on previous studies^13,25^.

The vaccination assay was done by using combination treated EMT6.huHER2 cells. Briefly, cells were treated with T-DM1 (5 ug/ml), BO-264 (500 nM) or their combination for 72 hours followed by collecting dying cells, washing twice with PBS and injection into the flank region of BALB/c mice with 5-7 mice per group. After a week, 1 million alive EMT6.huHER2 cells were injected into the opposite flank. Tumor formation was monitored, and % of tumor-free mice was recorded.

### Bioinformatics and statistical analyses

The microarray data set, GSE194040^26^ was download from the GEO database^27^. The ICD gene signature was retrieved from^28^. The expression levels of the genes in the T-DM1+pertuzumab-treated patients from GSE194040 were represented as a heat map using Morpheus software from Broad Institute, https://software.broadinstitute.org/morpheus. The significance between the number of responders vs. non-responders among low vs high ICD score expressers were determined using Chi-square testing. The significance between TACC3 protein expression in responder vs. non-responder patients upon T-DM1 treatment from the Hacettepe cohort was calculated using one sample Wilcoxon signed-rank test. All the results are represented as mean□±□standard deviation (SD) or mean□±□standard error of the mean (SEM), as indicated in the figure legends. All statistical analyses were performed in GraphPad Prism Software (v10.0.0 (153)) (San Diego, CA, USA). Comparisons between two groups were done using unpaired two-sided Student’s t-test. Tumor volumes between combination groups vs. single agent or vehicle-treated groups were compared using two-way ANOVA with the integration of Dunnett’s multiple comparison test. Survival curves for the vaccination assay were generated using Kaplan-Meier method, and significance between groups was calculated by Log-rank test. Experiments were repeated two to three times independently with similar results.

All other methods, including Western blotting, inhibitor treatments, CRISPR/Cas9-mediated gene knockout and stable overexpression, immunofluorescence staining, multiplex IHC staining, and multiplex cytokine array are provided in Supplementary Methods.

### Ethical approval

The use of human tissues from Hacettepe University was approved by the Non□Interventional Clinical Research Ethics Committee of Hacettepe University (approval no: 2020/02-40). The animal experiments were approved by the Institutional Animal Care and Use Committee of the University of South Carolina and Medical University of South Carolina.

### Data availability

Gene expression data were downloaded from the NCBI Gene Expression Omnibus database under GSE194040^26^.

## Results

### T-DM1 induces ICD markers in T-DM1 sensitive breast cancer cells in a mitotic arrest-dependent manner and ICD correlates with T-DM1 sensitivity in patients

To test if immunogenic cell death (ICD) is a mechanism of T-DM1 sensitivity, we first analyzed the phosphorylation of eIF2α, one of the major hallmarks of ICD^29^ upon T-DM1 treatment, and observed a prominent increase in p-eIF2α (S51) along with microtubule disassembly, mitotic arrest and apoptosis in two different HER2+ cell lines, SK-BR-3 and BT-474 (**Fig. 1A and Supplementary Fig. S1A**). This prompted us to test if T-DM1 would activate also other ICD-related DAMPs that are indispensable for induction of ICD. We observed that upon T-DM1 treatment, there was a significant increase in ATP secretion (**Fig. 1B**), HMGB1 release (**Fig. 1C**) and calreticulin cell surface exposure (**Fig. 1D, Supplementary Fig. S1B**) in both cell lines. We then assessed the cytokine profiles secreted from sensitive cells upon T-DM1 treatment and observed significant induction of IFN-gamma, IL12-p40, RANTES (CCL-5), IL-2 IL-13, IL-15, MCP-1 (CCL-2) and MCP-2 (CCL-7) (**Fig. 1E and Supplementary Fig. S1C**), which are well-established cytokines involved in ICD^30–33^. These results show that T-DM1-induced mitotic arrest and apoptosis are accompanied by increased eIF2α phosphorylation, induction of DAMPs and cytokines, indicative of ICD induction.

**Figure 1.**
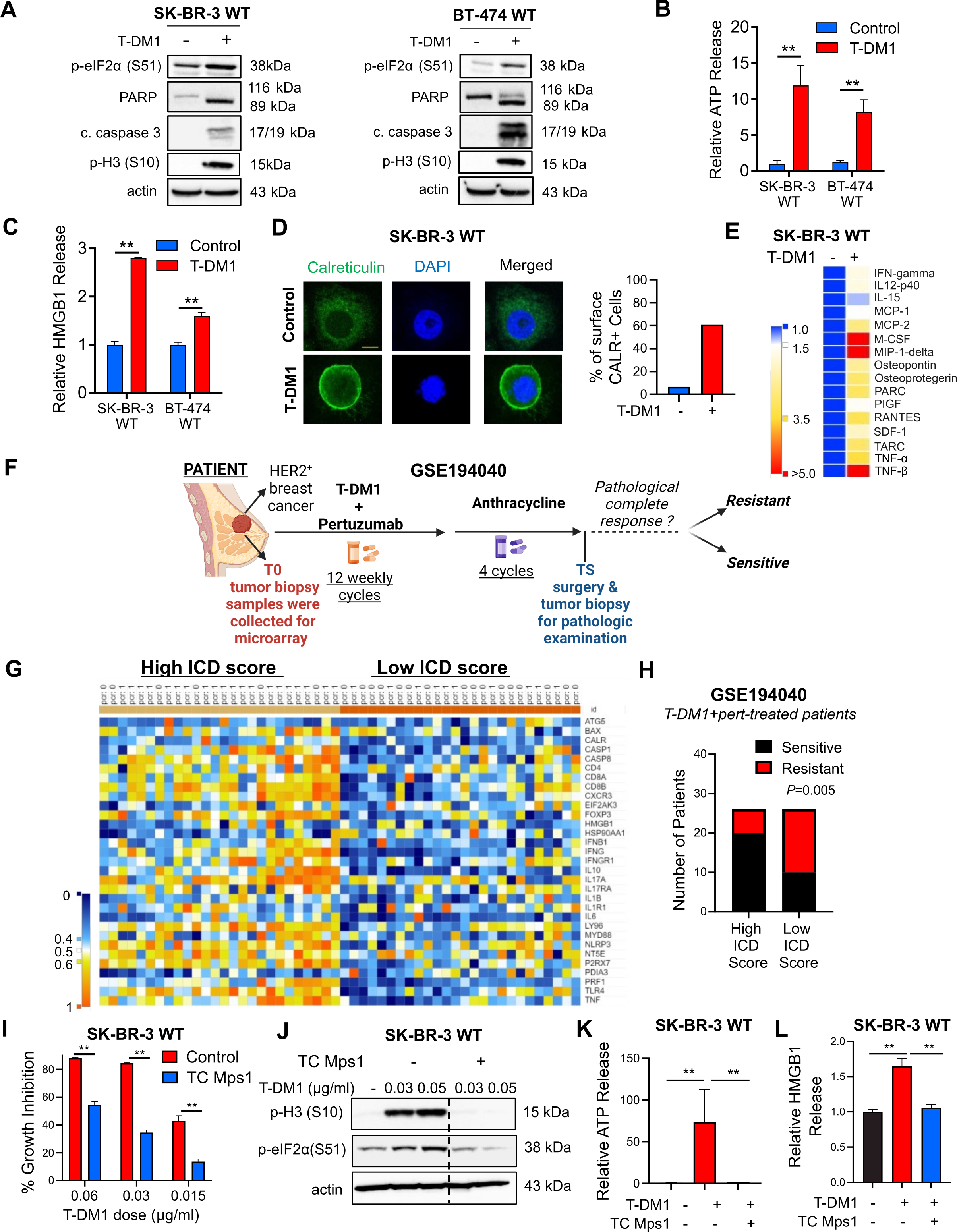
T-DM1 induces ICD markers in T-DM1 sensitive breast cancer cells in a mitotic arrest-dependent manner and ICD correlates with T-DM1 sensitivity in patients. **A** Western blot analysis of mitotic arrest, apoptosis and ICD markers in T-DM1-treated SK-BR-3 WT (left) and BT-474 WT (right) cells. **B, C** Relative ATP release (B) and HMGB1 release (C) from T-DM1-treated SK-BR-3 WT and BT-474 WT cells (n=3, 4). **D** IF cell surface staining of calreticulin (green) in T-DM1 treated SK-BR-3 WT cells. Scale bar=10 µm. DAPI was used to stain the nucleus. Its quantification is provided on the right. **E** Cytokine array blot analysis showing the differentially secreted cytokines in T-DM1-treated SK-BR-3 WT cells. **F.** Schematic summary of the treatment scheme and the sample collection timeline in GSE194040^26^. This figure was drawn using Biorender.com. **G.** Heatmap of ICD-related genes found in the ICD gene signature score^28^ and their correlation with pCR in T-DM1+pertuzumab-treated patients from GSE194040. pCR: 1, sensitive; pCR: 0, resistant. **H**. Chi-square analysis of sensitive vs. resistant tumors expressing low vs. high ICD score from G. **I** Percent growth inhibition in SK-BR-3 WT cells treated with T-DM1 alone or in combination with 1 µM TC Mps1 (Mps1 inhibitor) (n=4). **J** Western blot analysis of p-H3 and p-eIF2α in SK-BR-3 WT cells treated with T-DM1 alone or in combination with 1 µM TC Mps1. Actin is used as a loading control. **K, L** Relative ATP (K) and HMGB1 (L) release in SK-BR-3 WT cells treated with T-DM1 alone or in combination with 1 µM TC Mps1 (n=3). Data correspond to mean values□±□standard deviation (SD). *P*-values for the bar graphs were calculated with the unpaired, two-tailed Student’s t test. Significance for the Chi-square analysis was calculated with Chi-square testing. **, *P*<0.01.

To further test the clinical relevance of ICD in determining response to T-DM1, we analyzed the GSE194040^26^ patient dataset, containing gene expression profiling data from pre-treated HER2+ tumors of patients with clinical information on the status of pathologic complete response (pCR) or non-pCR after T-DM1+pertuzumab therapy **(Fig. 1F**). We generated a heatmap of genes found in an ICD-related gene signature^28^ (**Fig. 1G**) and correlated its levels with pCR. We showed that genes of the ICD score were expressed at higher levels in sensitive patients, while resistant patients were enriched in low ICD score expressers (**Fig. 1H**). These data suggest that ICD is a novel and clinically relevant mechanism of action of T-DM1 that correlates with clinical T-DM1 response.

To test if the T-DM1-induced release of DAMPs is dependent on mitotic arrest and activation of SAC, which are known to induce apoptosis under T-DM1 treatment^11,12^, we inhibited Mps1, one of the major regulators of SAC activation^34^, with TC Mps1 in T-DM1-treated SK-BR-3 cells. We observed that T-DM1 induced growth inhibition (**Fig. 1I** and **Supplementary Fig. S1D**), eIF2α phosphorylation (**Fig. 1J**), ATP release **(Fig. 1K**), HMGB1 release **(Fig. 1L, Supplementary Fig. S1E**) as well as cell surface exposure of calreticulin (**Supplementary Fig. S1F**) were all reversed upon SAC inhibition. To test if the observed induction of ICD markers is due to the antibody or the payload component of T-DM1, we tested the effect of trastuzumab, the antibody component of T-DM1, and showed that trastuzumab used at the same dose and duration as T-DM1 was not able to elicit the induction of eIF2α phosphorylation, ATP release or calreticulin surface exposure **(Supplementary Fig. S2A-D**), suggesting that T-DM1 induced ICD markers are due to its payload. Overall, our data show that T-DM1 induces mitotic cell death and activation of ICD markers in a SAC-dependent manner.

### ICD-related factors are lost in T-DM1 resistance upon TACC3 overexpression, TACC3 correlates with clinical T-DM1 resistance, and its inhibition overcomes T-DM1 resistance and restores ICD markers in vitro

It is unclear if the loss of ICD is a mechanism of drug resistance. Therefore, we first examined the markers of mitotic arrest, apoptosis and ICD in our previously published acquired resistant models of T-DM1 (BT-474 T-DM1R and SK-BR-3 T-DM1R)^9^. We observed that while T-DM1 induced mitotic arrest, apoptosis and eIF2α phosphorylation in wild-type (WT) (i.e., sensitive) cells, none of these markers were induced in T-DM1R cells (**Fig. 2A**). Supporting these data, T-DM1 was not able to elicit the induction of DAMPs (e.g., ATP release and HMGB1 release) in T-DM1R cell lines (**Supplementary Fig. S3A-B**). To identify the mediators of T-DM1 resistance that suppress the prolonged mitosis and ICD-related markers, which are induced in sensitive cells upon T-DM1 treatment, we examined the changes in mitosis genes upon T-DM1 resistance by re-analyzing our RNA-Seq data of BT-474 T-DM1R and SK-BR-3 T-DM1R cell lines compared to their WT counterparts (**Fig. 2B**). 21 genes were differentially expressed only in BT-474 T-DM1R cells while 35 genes were differentially expressed only in SK-BR-3 T-DM1R cells. 11 genes were deregulated in both cell lines in the same direction, among which TACC3 was significantly upregulated in the T-DM1R versions of both cell lines (**Fig. 2B**). We validated the increased expression of TACC3 in T-DM1 resistant cells also at protein level (**Fig. 2C**). To further test the clinical relevance of TACC3 protein expression in terms of its correlation with T-DM1 response, we stained TACC3 protein in tumor samples of HER2+ breast cancer patients with variable clinical outcome, having been collected before and after T-DM1 therapy in neo-adjuvant settings. While patients who responded to T-DM1 treatment have reduced TACC3 protein during T-DM1 treatment, the non-responders exhibited increased TACC3 expression over time (**Fig. 2D, E**), validating the association of TACC3 expression in clinical resistance to T-DM1.

**Figure 2.**
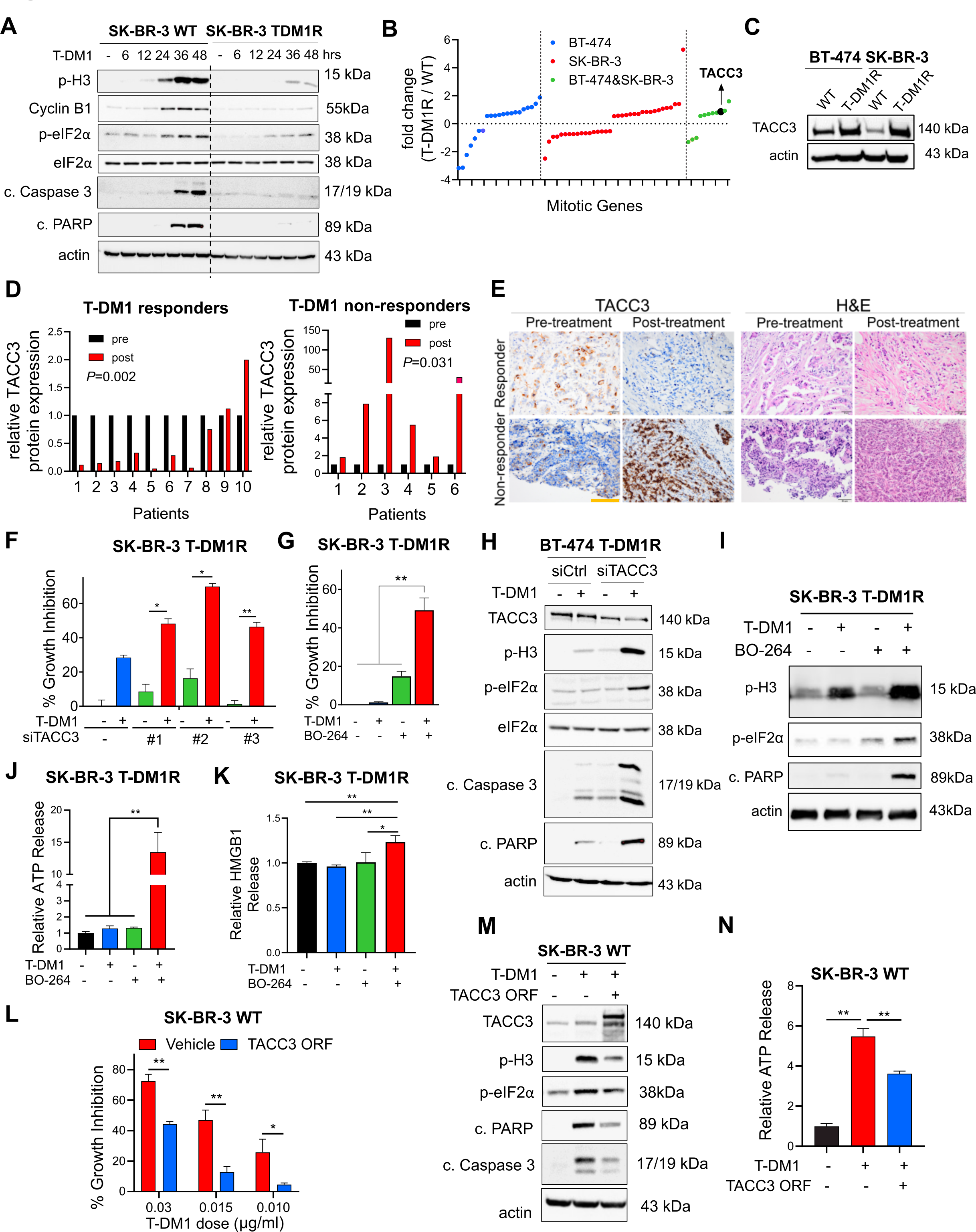
ICD-related factors are lost in T-DM1 resistance upon TACC3 overexpression, TACC3 correlates with clinical T-DM1 resistance, and its inhibition overcomes T-DM1 resistance and restores ICD markers in vitro. **A** Western blot analysis of mitotic arrest, apoptosis, and ICD markers in SK-BR-3 WT and T-DM1 resistant (T-DM1R) cells treated with 0.05 µg/mL T-DM1 in a time-dependent manner. **B** The log fold change of the mitotic genes differentially expressed only in BT-474, SK-BR-3 or both T-DM1R cells compared to WT counterparts in RNA-seq analysis. **C** Western blot analysis of TACC3 protein expression in BT-474 and SK-BR-3 WT vs. T-DM1R cells. Actin is used as a loading control. **D** Bar graphs showing relative protein expression of TACC3 in the T-DM1-treated tumors of responder vs. non-responder patients before and after treatment (n=6-10). **E** Representative TACC3 IHC and H&E staining in the tumor tissues of patients from D. Scale bar=100 µm. **F** Percent growth inhibition in SK-BR-3 T-DM1R cells transfected with siTACC3 and treated with 0.03 µM T-DM1 (n=4-6). **G** Percent growth inhibition in SK-BR-3 T-DM1R cells treated with T-DM1 alone or in combination with 1 µM TACC3 inhibitor (BO-264) (n=4-6). **H** Western blot analysis of mitotic arrest, apoptosis, and ICD markers in BT-474 T-DM1R cells transfected with siTACC3 and treated with T-DM1. Actin is used as a loading control. **I** Western blot analysis of mitotic arrest, apoptosis, and ICD markers in SK-BR-3 T-DM1R cells treated with T-DM1 alone or in combination with BO-264. Actin is used as a loading control. **J** Relative ATP release from SK-BR-3 T-DM1R cells treated with T-DM1 alone or in combination with BO-264 (n=3, 4). **K** Relative HMGB1 release from SK-BR-3 T-DM1R cells treated with T-DM1 alone or in combination with BO-264 (n=3). **L** Percent growth inhibition in SK-BR-3 WT cells overexpressing TACC3 and treated with T-DM1 (n=3). **M** Western blot analysis of mitotic arrest, apoptosis and ICD markers in SK-BR-3 WT cells overexpressing TACC3 and treated with T-DM1. Actin is used as a loading control. **N** Relative ATP release from SK-BR-3 WT cells overexpressing TACC3 and treated with T-DM1 (n=3). Data correspond to mean values□±□standard deviation (SD). Significance for D was calculated with one way Wilcoxon signed-rank test. *P*-values for other subfigures were calculated with the unpaired, two-tailed Student’s t test. *, *P*<0.05; **, *P*<0.01.

To determine the causal role of TACC3 overexpression in mediating T-DM1 resistance, we tested if targeting TACC3 overcomes T-DM1 resistance in our acquired T-DM1 resistant models. Inhibition of TACC3 either with three different siRNAs (**Fig. 2F, Supplementary Fig. S4A**) or with pharmacological inhibitors BO-264^13^ (**Fig. 2G, Supplementary Fig. S4B**) or SPL-B^35^ (**Supplementary Fig. S4C**) overcame resistance in both T-DM1R cell lines. TACC3 inhibition in combination with T-DM1 disrupted microtubule dynamics (**Supplementary Fig. S4D**), inhibited microtubule polymerization (**Supplementary Fig. S4E**) and caused mitotic arrest and apoptosis (**Fig. 2H, I, Supplementary Fig. S4F**). Notably inhibiting the SAC kinase, Mps1 using TC Mps1 reversed the TACC3 inhibition-mediated T-DM1 sensitization (**Supplementary Fig. S4G**), suggesting that SAC activation is also crucial for restoration of T-DM1 sensitivity by TACC3 targeting. Furthermore, combination treatment increased eIF2α phosphorylation (**Fig. 2H, I, Supplementary Fig. S4F**), ATP release (**Fig. 2J, Supplementary Fig. S5A-C**), HMGB1 release (**Fig. 2K, Supplementary Fig. S5D**) and calreticulin surface exposure in these T-DM1R cell lines (**Supplementary Fig. S5E-K**). It also induced secretion of pro-inflammatory cytokines (**Supplementary Fig. S5L**). Notably, overexpression of TACC3 in T-DM1 sensitive cells conferred T-DM1 resistance (**Fig. 2L**), abrogated T-DM1 induced mitotic arrest, apoptosis, eIF2α phosphorylation (**Fig. 2M**) and decreased ATP release (**Fig. 2N**). Overall, our data show that T-DM1 resistance is characterized by loss of ICD markers, and targeting TACC3 overcomes T-DM1 resistance in a SAC-dependent manner and restores the induction of T-DM1-induced ICD markers in vitro.

### Targeting TACC3 in combination with T-DM1 in human HER2-expressing murine cells induces ICD markers and leads to ex vivo DC maturation and T cell activation

To test the effects of TACC3 inhibition in combination with T-DM1 in a syngeneic T-DM1 resistant setting that will allow us to assess the changes in the immunogenicity of the cells, we first developed the human HER2-overexpressing derivative of the murine EMT6 mammary tumor cells; EMT6.huHER2. We demonstrated that these cells are resistant to T-DM1, in line with the literature^36^, and inhibiting TACC3 with BO-264 reversed resistance by reducing cell viability in a dose-dependent manner (**Fig. 3A**). Furthermore, we validated our results by knocking out TACC3 using CRISPR/Cas9 system. Both sgTACC3 constructs effectively reduced TACC3 expression (**Fig. 3B**) and mediated T-DM1 sensitization (**Fig. 3C**). Notably, combination of TACC3 knockout or its pharmacologic inhibition with T-DM1 induced mitotic arrest, eIF2α phosphorylation, and apoptosis (**Fig. 3D, E** and **Supplementary Fig. S6**) and activated the ICD markers; ATP release (**Fig. 3F, G**) and calreticulin cell surface exposure (**Fig. 3H, I**) in the T-DM1 resistant EMT6.huHER2 cells.

**Figure 3.**
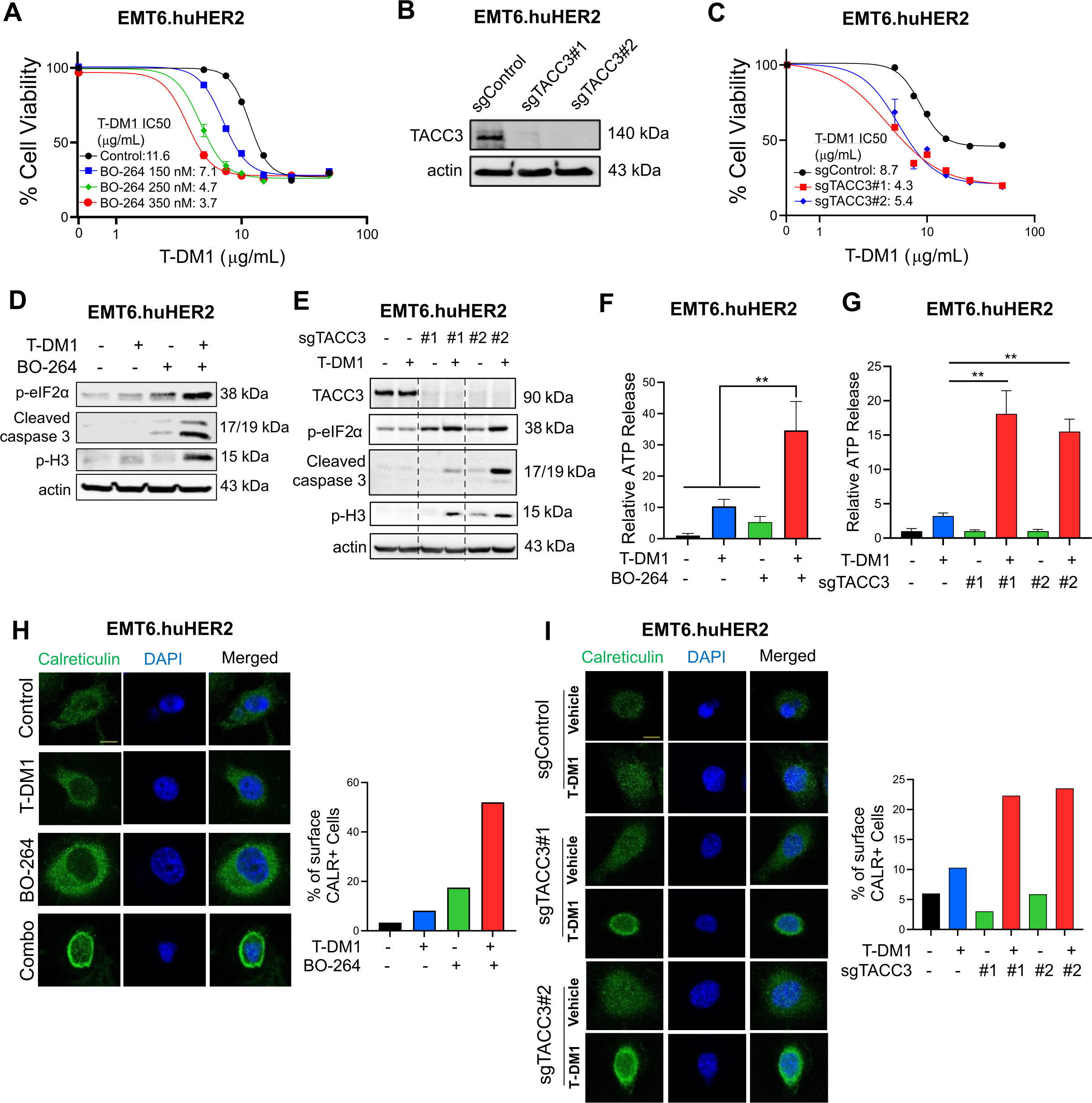
Targeting TACC3 sensitizes the human HER2-expressing EMT6.huHER2 cells to T-DM1 and induces ICD markers. **A** Cell viability assay in EMT6.huHER2 cells treated with increasing doses of T-DM1 alone or combination with different dose of BO-264 for 3 days (n=4). **B** Validation of TACC3 knockout in EMT6.huHER2 cells obtained using CRISPR/Cas9 system. **C** Cell viability assay in EMT6.huHER2.sgTACC3 vs. sgControl cells treated with increasing doses of T-DM1 for 3 days (n=4). **D** Western blot analysis of mitotic arrest, apoptosis and ICD markers in EMT6.huHER2 cells treated with T-DM1 alone or in combination with BO-264. Actin is used as a loading control. **E**. Western blot analysis of TACC3, mitotic arrest, apoptosis, and ICD markers in EMT6.huHER2.sgTACC3 vs. sgControl cells treated with T-DM1. Actin is used as a loading control. **F** Relative ATP release from EMT6.huHER2 cells treated with T-DM1 alone or in combination with BO-264 (n=3). **G** Relative ATP release from EMT6.huHER2.sgTACC3 vs. sgControl cells treated with T-DM1 (n=3, 4). **H** IF cell surface staining of calreticulin (green) in EMT6.huHER2 cells treated with T-DM1 alone or in combination with BO-264. Its quantification is provided on the right. **I** IF cell surface staining of calreticulin (green) in EMT6.huHER2.sgTACC3 vs. sgControl cells treated with T-DM1. Its quantification is provided on the right. Data correspond to mean values□±□standard deviation (SD). *P*-values were calculated with the unpaired, two-tailed Student’s t test. **, *P*<0.01.

It has been shown that ICD induction is followed by DC maturation that further results in T cell activation^37,38^. To test whether the combination of TACC3 inhibition with T-DM1 induces DC maturation and T cell activation, we isolated bone marrow-derived DCs from BALB/c mice (the strain EMT6 cells were originated from) (**Fig. 4A**). Incubation of DCs with conditioned media (CM) collected from combination treated EMT6.huHER2 cells resulted in increased expression of the DC maturation markers, CD80 and CD86 (**Fig. 4B, D, Supplementary Fig. S7A**). Furthermore, co-culturing these maturated DCs with T cells increased T cell activation as shown by CD8 and CD25 staining (**Fig. 4E, G, Supplementary Fig. S7B**). These results were also recapitulated using CRISPR-Cas9-mediated knockout of TACC3 (**Fig. 4C, D, F, G**). We profiled the secreted cytokines/chemokines using multiplex cytokine analysis, and observed that proinflammatory markers IL-1β, IL-2, IL-6, IL-17, MIP-1α, MIP-1β and TNF-α were all increased in combination treated samples (**Fig. 4H**). Overall, these data suggest that inhibition of TACC3 in combination with T-DM1 induces ICD markers and leads to ex vivo DC maturation and T cell activation in EMT6.huHER2 murine mammary tumor model.

**Figure 4.**
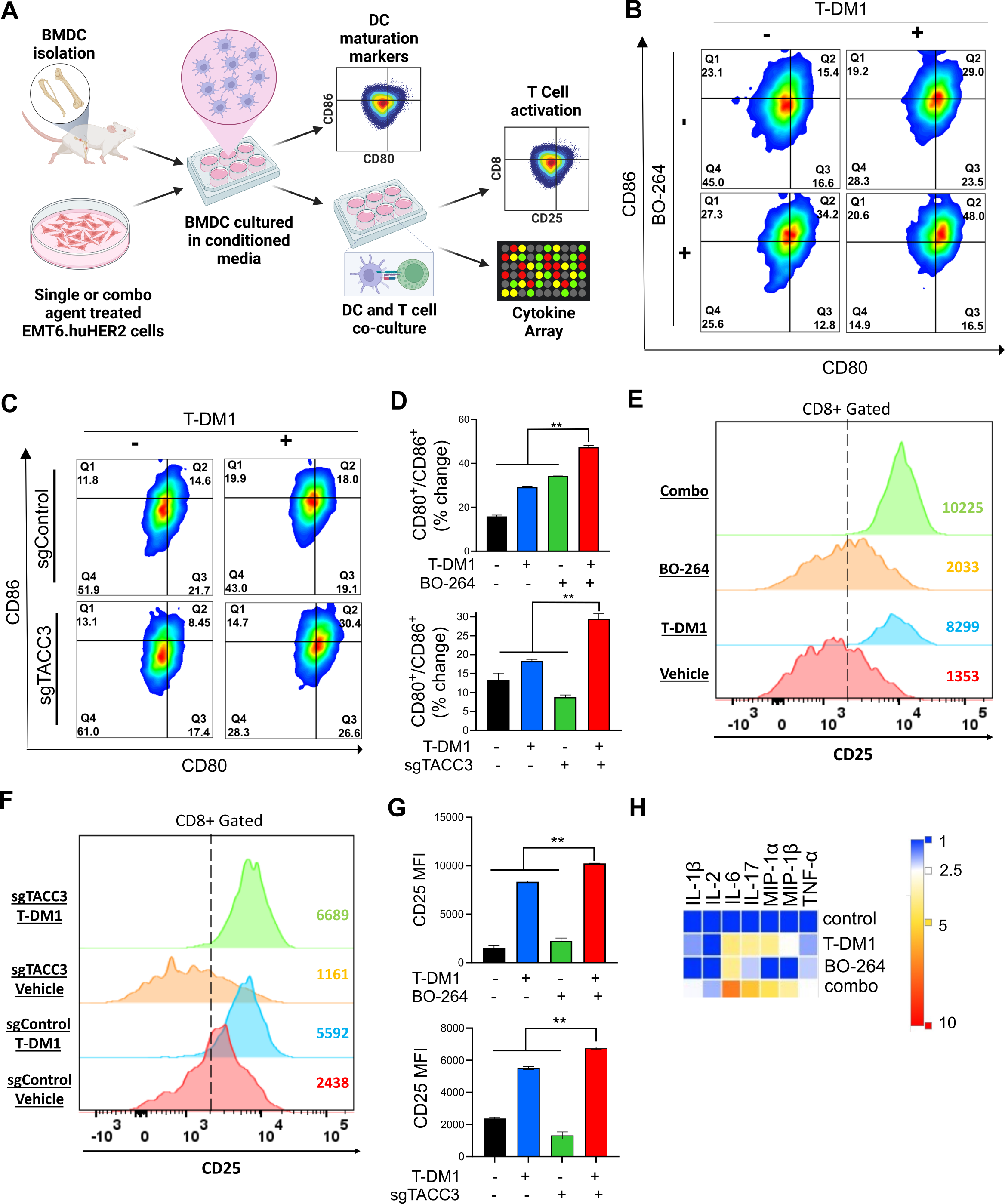
Inhibition of TACC3 in combination with T-DM1 leads to ex vivo DC maturation, T cell activation, and release of ICD related pro-inflammatory cytokines. **A** Schematic representation of the experimental workflow for DC maturation, T cell activation and cytokine profiling experiments, drawn using Biorender.com. **B, C** Flow cytometry analysis of DC maturation markers in DC cells incubated with the conditioned media collected from EMT6.huHER2 cells treated with 7.5 µg/ml T-DM1 and 500 nM BO-264, alone or in combination (B) or in EMT6.huHER2.sgControl vs. sgTACC3 cells treated with 7.5 µg/ml T-DM1 (C). **D** Quantification of CD80+/CD86+ cells from B and C (n=2). **E, F** Flow cytometry analysis of T cell activation marker, CD25 in CD8+ T cells co-cultured with DCs from B and C. **G** Quantification of the CD25 mean fluorescence intensity (MFI) from E and F (n=2). **H** Levels of pro-inflammatory cytokines in the media collected from DC-T cell co-cultures from E, F. Data correspond to mean values□±□standard deviation (SD). *P*-values were calculated with the unpaired, two-tailed Student’s t test. **, *P*<0.01.

### Targeting TACC3 induces ICD, leads to immune cell infiltration and potentiates T-DM1 response in vivo

To assess the effects of the combination of T-DM1 with TACC3 inhibition on ICD induction in vivo, we performed the so-called vaccination assay using the T-DM1 resistant EMT6.huHER2 cells^39^. Cells were treated with T-DM1 or BO-264 alone or in combination for 72 hours, and the dying cells were injected into the flank of BALB/c mice, followed by injection of living cells to the opposite flank after a week, and monitoring of tumor formation (**Fig. 5A**). Vaccinating mice with dying cells treated with our combination therapy significantly improved tumor-free survival compared to control or single-agent treated groups, demonstrating that our combination therapy elicits ICD in vivo to inhibit tumor formation (**Fig. 5B**).

**Figure 5.**
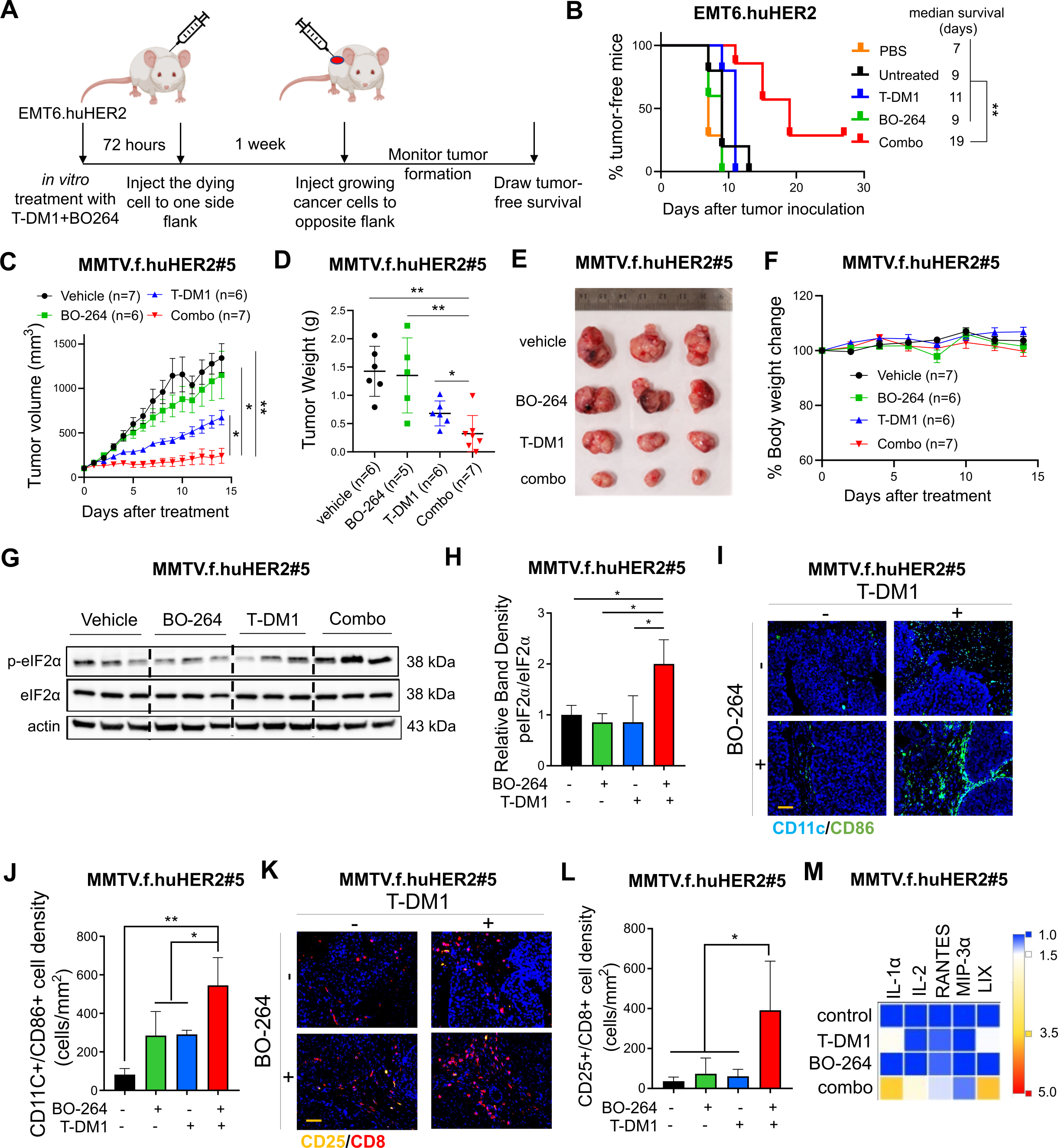
TACC3 inhibition elicits ICD in vivo and potentiates TDM1 response via increasing the infiltration of anti-tumor immune cells in vivo. **A** Schematic representation of the in vivo vaccination assay drawn using Biorender.com. **B** Tumor-free survival curves of BALB/c mice vaccinated with PBS or single agent or combination treated EMT6.huHER2 cells (n=5-7). **C** Tumor growth of the MMTV.f.huHER2#5 model under low dose T-DM1 (5 mg/kg, once) in combination with BO-264 (50 mg/kg, daily) (n=6, 7). **D** Tumor weights of the mice in C after 14 days of treatment. **E, F** Representative resected tumor pictures (E) and body weights (F) from mice in C. **G** Western blot analysis of p-eIF2α and eIF2α protein expression levels in tumors from C. Actin is used as a loading control. **H** Relative band density graphs for p-eIF2α normalized to eIF2α from G (n=3). **I, J** Multiplex IF staining of CD11c/CD86 and CD25/CD8 in short-term-treated MMTV.f.huHER2#5 tumors and its quantification (n=3). **K, L** Multiplex IF staining of CD11c/CD86 and CD25/CD8 in short-term-treated MMTV.f.huHER2#5 tumors and its quantification (n=3). Scale bar=100 µm. **M** Levels of the cytokines in the serums of the mice with short-term-treated MMTV.f.huHER2#5 tumors (n=3). Data for the bar graphs and box plots correspond to mean values□±SD, while data for the tumor volume and body weight graphs correspond to mean values□± standard error of the mean (SEM). End-point criteria for mice in C and F are treatment for 14 days or until reaching ethical tumor size cut-off. *P*-values for the bar graphs and box plots were calculated with the unpaired, two-tailed Student’s t test. Significance for the tumor volume graph and multiplex IHC quantification was calculated with two-way and one-way ANOVA, respectively. *, *P*<0.05; **, *P*<0.01.

To test the effects of TACC3 inhibition on enhancing T-DM1 response in the first-line settings, we utilized the relatively T-DM1 responsive human HER2-overexpressing MMTV.f.huHER2#5 (Fo5) transgenic model^8,40^. Combination treatment completely blocked tumor growth as compared to single agent T-DM1 (5 mg/kg, i.v., once) or BO-264 (50 mg/kg, oral gavage, daily) treatments (**Fig. 5C-E**), without affecting body weight (**Fig. 5F**). Importantly, combination-treated Fo5 tumors exhibited the highest levels of p-eIF2α, the canonical marker of ICD (**Fig. 5G, H**). To analyze changes in tumor infiltrated lymphocytes (TIL) and tumor infiltrated DCs (TIDCs) upon combination treatment, we treated the Fo5 tumor-bearing mice with T-DM1 alone or in combination with BO-264 for a week and performed multiplex IHC staining of DCs and T cells in the collected tumors. As a result, we observed that the infiltration of CD11c+CD86+ mature DCs (**Fig. 5I, J**) and CD8+CD25+ activated cytotoxic T cells (**Fig. 5K, L**) were significantly increased in tumor samples treated with the combination therapy. Notably, there was a trend towards a decrease in T-DM1-induced infiltration of tumor promoting Foxp3+/CD4+ regulatory T (Treg) cells, while NK1.1+/CD27+ NK cells that are responsible for trastuzumab-mediated antibody dependent cellular cytotoxicity (ADCC), underwent a slight, albeit not significant, decrease in combination therapy (**Supplementary Fig. S8**). We further analyzed the serum levels of cytokines collected from treated mice and observed an increase in the pro-inflammatory cytokines (IL-1α, IL-2, RANTES (CCL5), MIP-3α and LIX (CXCL5)) (**Fig. 5M**). Altogether, these data demonstrate that TACC3 inhibition restores T-DM1-induced ICD and increases the infiltration of anti-tumor DCs and cytotoxic T cells without triggering the infiltration of pro-tumorigenic Tregs, thus leading to stronger growth inhibition of human HER2-expressing tumors.

## Discussion

ICD is a unique form of cell death that can activate anti-tumor immune response and has been shown to be induced by several different anti-cancer therapies. Despite being the first and one of the most successful antibody-drug conjugates (ADCs) approved for refractory HER2-positive breast cancer, little is known if response and resistance to trastuzumab emtansine (T-DM1) involves ICD modulation that can be leveraged to enhance T-DM1 response. Here, we demonstrate, for the first time, that the iconic ADC, T-DM1 can elicit all the hallmarks of ICD, i.e., eIF2α phosphorylation, ATP secretion, HMGB1 release and calreticulin surface exposure in a SAC-dependent manner in drug sensitive models (**Fig. 6A**). In T-DM1 resistance, TACC3 is upregulated and inhibits T-DM1-induced SAC activation, thus blocking eIF2α phosphorylation and DAMPs, which ultimately results in cell survival **(Fig. 6B**). Inhibiting TACC3 in T-DM1 resistant tumors restores T-DM1-induced SAC activation, mitotic cell death, eIF2α phosphorylation and elevated levels of DAMPs, i.e., the hallmarks of ICD. TACC3 inhibition in combination with T-DM1 further induces pro-inflammatory cytokine secretion, DC maturation and T cell activation, eventually causing inhibition of tumor growth (**Fig. 6C**).

**Figure 6.**
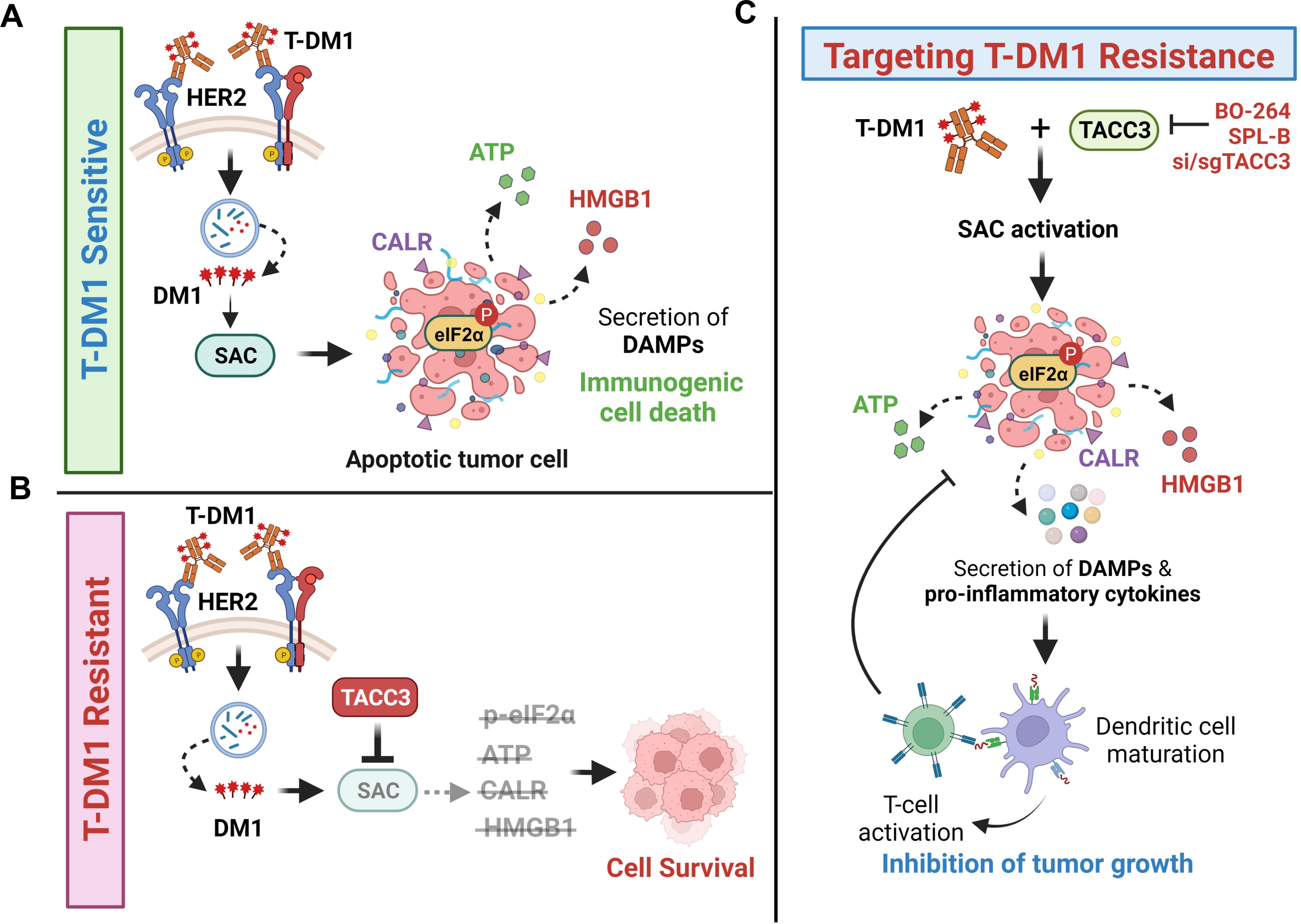
Schematic summary of the proposed model of T-DM1 sensitivity, resistance and targeting T-DM1 resistance. **A** In T-DM1 sensitive tumors, the activation of spindle assembly checkpoint (SAC) and mitotic arrest lead to apoptosis and activation of ICD markers, e.g., eIF2α phosphorylation, ATP secretion, calreticulin surface exposure, and HMGB1 release, leading to DC maturation and cytotoxic T cell, culminating in tumor growth inhibition. **B** In T-DM1 resistant tumors, overexpression of TACC3 prevents activation of SAC, mitotic cell death and ICD, thus promoting cell survival. **C** Inhibition of TACC3 in combination with T-DM1 in the resistant tumors restores SAC activation and mitotic arrest, leading to apoptosis and ICD induction, thereby increasing the infiltration of DCs and T cells, thus restoring T-DM1 sensitivity. This figure was drawn using Biorender.com.

In recent years, it has been increasingly recognized that anti-tumor immune activation is an integral part of response to anti-cancer therapies. To evoke a cytotoxic immune response, these therapies trigger an immunogenic form of cell death that is characterized by secretion of the so called ‘eat me’ signals, i.e., DAMPs from the dying cancer cells. These DAMPs then lead to infiltration and activation of anti-tumor immune cells, leading to tumor shrinkage. Along these lines, a higher level of tumor infiltrating lymphocytes (TILs) usually indicates better response to therapy^41^. Recently, an open-label, phase III study (KRISTINE) has reported an association of TIL infiltration with T-DM1 response^42^. In T-DM1+pertuzumab-treated patients, those expressing higher HER2 and immune marker levels exhibited higher pCR rates^42^, suggesting a potential involvement of anti-tumor immune activation in T-DM1 sensitivity in the patients. Association of TILs with clinical response has also been tested for other ADCs, e.g., trastuzumab deruxtecan and a positive correlation between number of TILs and drug sensitivity was observed^43,44^. However, it has yet to be determined whether the potential link between the clinical response to T-DM1 or other ADCs and TILs is molecularly mediated by elicitation of ICD. Here, we demonstrated, for the first time, that ICD induction is one of the mechanisms of T-DM1 sensitivity in vitro and in vivo. Importantly, we showed that an ICD-related gene signature positively correlates with response to T-DM1 containing therapies in patients, suggesting that the observed association between TILs and T-DM1 sensitivity in the clinic may, in part, stem from ICD induction. Therefore, our data encourages future clinical studies investigating the association between sensitivity to ADCs, including T-DM1 and ICD hallmarks.

Anti-microtubule agents, such as taxanes are among the most commonly used chemotherapeutics in cancer and their major mechanism of action involves mitotic cell death via disruption of microtubule dynamics. Intriguingly, recent studies have suggested that the clinical success of microtubule-targeting agents is not only a result of mitotic cell death but potentially involves novel mechanisms that lead to activation of a strong anti-tumor immune response^45–47^. For instance, paclitaxel, one of the most widely used taxanes was shown to promote a proinflammatory response by activation of innate immunity^46,48^. Paclitaxel may also improve the efficacy of PD-L1 blockade therapy in animal models by causing tumor eradication, metastasis suppression, and preventing recurrence^49^. Furthermore, paclitaxel has recently been shown to activate ICD in ovarian cancer cells^50^. Despite these preliminary evidence on the potential roles of microtubule targeting agents on activating anti-tumor immunity, a mechanistic connection between mitotic arrest, SAC activation and the induction of ICD markers has not been tested before. In this study, we demonstrated that the T-DM1-induced mitotic arrest and SAC activation are required for the induction of ICD hallmarks, identifying SAC activation as a potentially novel way to induce ICD. However, given that ICD induction is, in part, dictated by the structure of the drug in addition to its molecular mechanisms of action, it has yet to be determined whether other inducers of SAC activation or mitotic arrest could also activate the hallmarks of ICD and elicit immune responses.

TACC3 is a microtubule and centrosome-associated protein playing key roles in mitotic progression^20,51^. TACC3 inhibition was shown to cause formation of multipolar spindles, mitotic arrest and apoptosis^13,21,35^. In recent years, novel non-canonical roles of TACC3 in tumor progression are also emerging. For instance, an *in silico* analysis in kidney renal cell carcinoma demonstrated that TACC3 expression correlates with several different types of immune cells, including follicular helper T cells and Tregs^52^. Infiltration of the immune suppressive Tregs into tumors may promote tumor growth via blocking anti-tumor immune responses^53^. For instance, a combination of T-DM1 with immune checkpoint blockers showed efficacy in animal models; however, the infiltration of tumor-promoting Foxp3+/CD4+ Tregs was shown to be increased as well^30^. Along these lines, although a trend is observed towards higher progression-free survival in PDL1+ patients upon combination of T-DM1 with the immune checkpoint inhibitor, atezolizumab, the difference in survival was not statistically meaningful with more adverse events^54^. These findings underlie the necessity to identify novel therapeutic strategies to achieve potent and durable immunogenic responses without activating tumor promoting immune subsets in larger patient subpopulations. Our data demonstrating the secretion of pro-inflammatory cytokines, DC maturation and increased infiltration of cytotoxic effector T cells upon inhibiting TACC3 in combination with T-DM1 without an increase in the infiltration of Foxp3+ Tregs suggest that TACC3 inhibition could be a superior therapeutic strategy to boost the immunogenicity of the tumors without activating tumor-promoting factors.

It has been demonstrated that T-DM1 treatment increases tumor infiltrating NK cells that are responsible for trastuzumab-induced ADCC^55,56^ in the absence or presence of immunotherapy^30^. Interestingly, we did not observe a significant change in the infiltration of NK cells upon T-DM1 treatment or upon treatment with the combination of T-DM1 and TACC3 inhibitor. The lack of NK cell infiltration even in T-DM1 monotherapy group is probably due to lower T-DM1 dose used in our study, i.e., 5 mg/kg compared to 15 mg/kg dose used in Muller et al^30^. Nonetheless, the strong tumor growth inhibition that we observed upon combination therapy with no NK cell infiltration suggest that TACC3 inhibition-mediated T-DM1 potentiation does not likely involve trastuzumab-mediated ADCC. Overall, our data encourages testing the combination of TACC3 inhibitors with other ADCs beyond T-DM1 or even with immune checkpoint blockers to achieve superior and durable responses.

Overall, we showed that ICD induction upon SAC-induced mitotic cell death is a novel mechanism of T-DM1 sensitivity and activates T cell-mediated anti-tumor immunity, while T-DM1 resistance is characterized by loss of ICD. We further identified TACC3 as a novel resistance mediator whose inhibition restores the induction of ICD hallmarks and increases the infiltration of cytotoxic effector T cells into tumors. These data provide preclinical evidence for targeting TACC3 to revive tumor immunogenicity driven by ICD-related DAMPs in T-DM1 refractory HER2+ breast cancer that may ultimately result in improved clinical outcome.

## Acknowledgements

We are thankful to the members of Ozgur Sahin laboratory for invaluable discussion and advice. We thank the Translational Science Laboratory and the Flow Cytometry & Cell Sorting Shared Resource of Medical University of South Carolina. We thank Dr. Stephen Royle (University of Warwick) for providing us TACC3 ORF-expressing vector.

## Declaration of Interest

O. Sahin, B.C. and E.B. are the co-founders of OncoCube Therapeutics LLC. O. Sahin is the president of LoxiGen, Inc. The other authors declare no potential conflicts of interest.

## Funding

This work was supported by research funding from Mary Kay Ash Foundation Grant MK-07-21 (O. S). and in part, from the National Institutes of Health (NIH, R01CA251374 to O.S.) and previously by TUBITAK-BMBF Bilateral Grants (TUBITAK, 214Z130 (OS) and BMBF WTZ, 01DL16003 (SW)). The core facilities utilized are supported by NIH (C06 RR015455), Hollings Cancer Center Support Grant (P30 CA138313), or Center of Biomedical Research Excellence (COBRE) in Lipidomics and Pathobiology (P30 GM103339). The Zeiss 880 microscope was funded by a Shared Instrumentation Grant (S10 OD018113).

## Author Contributions

M.E.G and Ozge S. designed and performed experiments, acquired and analyzed data, interpreted data, and prepared the paper; N.O. performed the DC maturation, co-culture and flow cytometry analysis, M.U. performed the TACC3 IHC staining and evaluation; O.A. performed cell-based assays; M.C. contributed to human cytokine array experiment; M. A. generated the EMT6.huHER2 cell line; K.I. synthesized the BO-264; B.C. contributed to BO-264 synthesis and data interpretation; E.B. contributed to BO-264 synthesis and data interpretation; S.W. performed RNA-seq experiments of the T-DM1R vs. WT cells and contributed to data analyses; A.U. performed the IHC staining of HER2+ patient tissues and contributed to data interpretation; S.A. provided clinical information of HER2+ patients and contributed to data analyses; S.M. contributed to the design of co-culture experiments, multiplex IHC staining and help with interpreting the results; O.S. designed the study, oversaw experiments and data analyses, and prepared the paper. All authors reviewed and commented on the paper.

