## Supplementary Information for "Reviving immunogenic cell death upon targeting TACC3 enhances T-DM1 response in HER2-positive breast cancer"

Ozgur Sahin, PhD

Professor and SmartState Endowed Chair

Department of Biochemistry and Molecular Biology

Hollings Cancer Center

Medical University of South Carolina

86 Jonathan Lucas Street, Room HO712F, Charleston, SC 29425

### Supplementary Methods

#### Western blotting

Protein isolation and Western blotting were done as previously described[1,2]. Briefly, RIPA buffer (150 mmol/L NaCl, 50 mmol/L Tris base pH 8.0, 1 mmol/L EDTA, 0.5% sodium deoxycholate, 1% NP40, 0.1% SDS, 1 mmol/L DTT, and 1 mmol/L Na3VO4) supplemented with protease and phosphatase inhibitor cocktails were used to isolate total protein lysate. Protein concentrations were measured using the BCA Protein Assay Reagent Kit (Thermo Fisher Scientific, MA, USA). Equal amounts of protein (15–20 μg) were separated using 10% or 12% SDS-PAGE gel. Separated proteins were transferred to PVDF membranes (Bio-Rad, CA, USA) using a Trans-Blot turbo transfer system (Bio-Rad, CA, USA) and incubated with primary antibodies listed in **Supplementary Table S2**. HRP-linked anti-mouse IgG (#7076) or anti-rabbit IgG antibodies (#7074) (Cell Signaling Technology, MA, USA) were used as secondary antibodies, and signals were detected by enhanced chemiluminescence (ECL) prime western blot detection reagent (Cytiva, MA, USA). Images were acquired using Image Lab Software (Biorad, CA, USA) or iBright Analysis Software (Thermo Fisher Scientific, MA, USA).

#### Inhibitor treatments

T-DM1 was obtained from Genentech under a material transfer agreement (OR-219728 and OR-224086A) and dissolved in 100% dd$H_2O$ to yield a stock concentration of 2 mg/ml. BO-264 (synthesized as reported previously[13]), TC Mps1 (Tocris Biosciences, Bristol, UK), SPL-B (Axon MedChem, VA, USA) were dissolved in 100% DMSO to yield a stock concentration of 10 mM. Trastuzumab was dissolved in 100% water to yield a stock concentration of 10 mg/ml. For cell viability assay, BT-474 WT and T-DM1R ($8x10^3$ cells/well), SK-BR-3 WT and T-DM1R ($6x10^3$ cells/well), EMT6.huHER2 ($3x10^3$ cells/well) cells were seeded, and inhibitor treatments were performed at different concentrations. Cell viability was measured 72 hours after treatment with SRB (Sigma Aldrich) as described previously[3]. Annexin V/PI staining (Biolegend, USA) was performed according to manufacturer's instructions using EMT6.huHER2 cells treated with 500 nM BO-264 and 5 µg/mL T-DM1, alone or in combination for 48 hours.

**Gene knockout via CRISPR/Cas9-mediated gene editing and stable overexpression**

To generate EMT6.huHER2 cells, the human HER2 sequence from the pLL-RSV-hHER2 vector that was kindly provided by Hasan Korkaya (Augusta University, GA, USA) was cloned into the pCDH-CMV vector. The sgRNAs targeting mouse TACC3 were designed and selected based on having high on-target (=high efficacy) and low off-target (=high specificity) activity using the CRISPick tool (Broad Institute). The sgRNA sequences targeting TACC3 in EMT6.huHER2 murine mammary tumor cell line are: 5'-CACCGAGTTTAAGGAGTCGGCCTGG-3' (sg#1) and 5'-CACCGCTGAGATCCTAAGAGCAGA-3' (sg#2). The designed sgRNAs were cloned into human lentiCRISPR v2 vector (Addgene, MA, USA). For lentiviral packaging, HEK293T cells were transfected with sgRNAs and the packaging plasmids, pMD2.G and psPAX2 (Addgene, MA, USA). Transduction of the cells was performed in the presence of 10 ug/ml polybrene, and selection of transduced cells was done using 1µg/ml puromycin. Subsequently, transduced cells were transferred to 96-well plates in 1:2 serially diluted for seeding single cell colonies[4].

**Immunofluorescence staining**

SK-BR-3 (150,000 cells /well) and BT-474 (300,000 cells/well) WT and T-DM1R and EMT6.huHER2 (270,000 cells/well) cells were plated on glass coverslip in 6-well cell culture plates and treated with 0.05, 4 µg/ml and doses 7.5 µg/ml of T-DM1 and or 500 nM BO-264 on the next day. Forty-eight hours after treatment, cells were fixed with 2% paraformaldehyde for 5 min. For examination of microtubule organization, cells were treated with T-DM1 for 24 hours. Later, they were blocked for 1 hour in blocking buffer (3%BSA in PBS) and incubated with primary antibodies (**Supplementary Table S2**) in blocking buffer for 1 hour at room temperature (RT). Antibodies against calreticulin was used at a dilution of 1:500, and alpha-tubulin at a dilution of 1:1000. Then, the cover slips were incubated with secondary antibodies in blocking buffer for 1 hour at RT in dark (1:1000 dilution). Cells were also counter stained with DAPI for 5 min (0.01 µg/µl). Finally, slides were mounted using Prolong™ Glass mounting (Invitrogen, MA, USA) and examined using Zeiss LSM 880 confocal laser scanning microscope.

**Multiplex IHC staining**

Immunofluorescence staining of FFPE tumor slides were performed by deparaffinization at 60 ℃ for 1 hour, followed by rehydration in citrisolv for 5 min (3 times), 100% ethanol for 5 min (twice), 95% ethanol for 5 min (twice), deionized water for 5 min (twice). Antigen retrieval was done with Tris-EDTA pH=9 at 96 ℃ for 15 min, followed by cooling down to room temperature for 30 min, and washing with TBST for 5 min. Blocking was done at room temperature with Buffer W (IBA-Lifesciences) for 30 min. CD86, CD11c, CD25 and CD8 immunofluorescence was employed using the OPAL multiplexing method based on Tyramide Signal Amplification (TSA) with Opal 520, 480, 620 and Opal 690, respectively. The entire samples were scanned using the Vectra® Polaris™ Automated Quantitative Pathology Imaging System (Akoya Biosciences). Segmentation of tissue, segmentation of cells and cell phenotyping were done using inForm® Tissue Analysis Software (v[2.6.0], Akoya Biosciences). Merging of data, consolidation and analysis were done using Rstudio v(2023.3.3.0+).

**Multiplex cytokine array**

The multiplex cytokine analyses were performed using (1) supernatants of the DC and T cell co-cultures, and (2) serum samples collected from Fo5 tumors of mice treated with T-DM1 alone or in combination with BO-264 for a week were performed using the Luminex™ 200 system (Luminex, Austin, TX, USA) by Eve Technologies Corp. (Calgary, Alberta, Canada). Forty-five markers were simultaneously measured in the samples using Eve Technologies' Mouse Cytokine 45-Plex Discovery Assay® which consists of two separate kits; one 32-plex and one 13-plex (MilliporeSigma, Burlington, Massachusetts, USA). The assay was run according to the manufacturer's protocol. The 32-plex consisted of Eotaxin, G-CSF, GM-CSF, IFNγ, IL-1α, IL-1β, IL-2, IL-3, IL-4, IL-5, IL-6, IL-7, IL-9, IL-10, IL-12(p40), IL-12(p70), IL-13, IL-15, IL-17, IP-10, KC, LIF, LIX, MCP-1, M-CSF, MIG, MIP-1α, MIP-1β, MIP-2, RANTES, TNFα, and VEGF. The 13-plex consisted of 6Ckine/Exodus2, Erythropoietin, Fractalkine, IFNβ-1, IL-11, IL-16, IL-20, MCP-5, MDC, MIP-3α, MIP-3β, TARC, and TIMP-1.

### Supplementary Figures

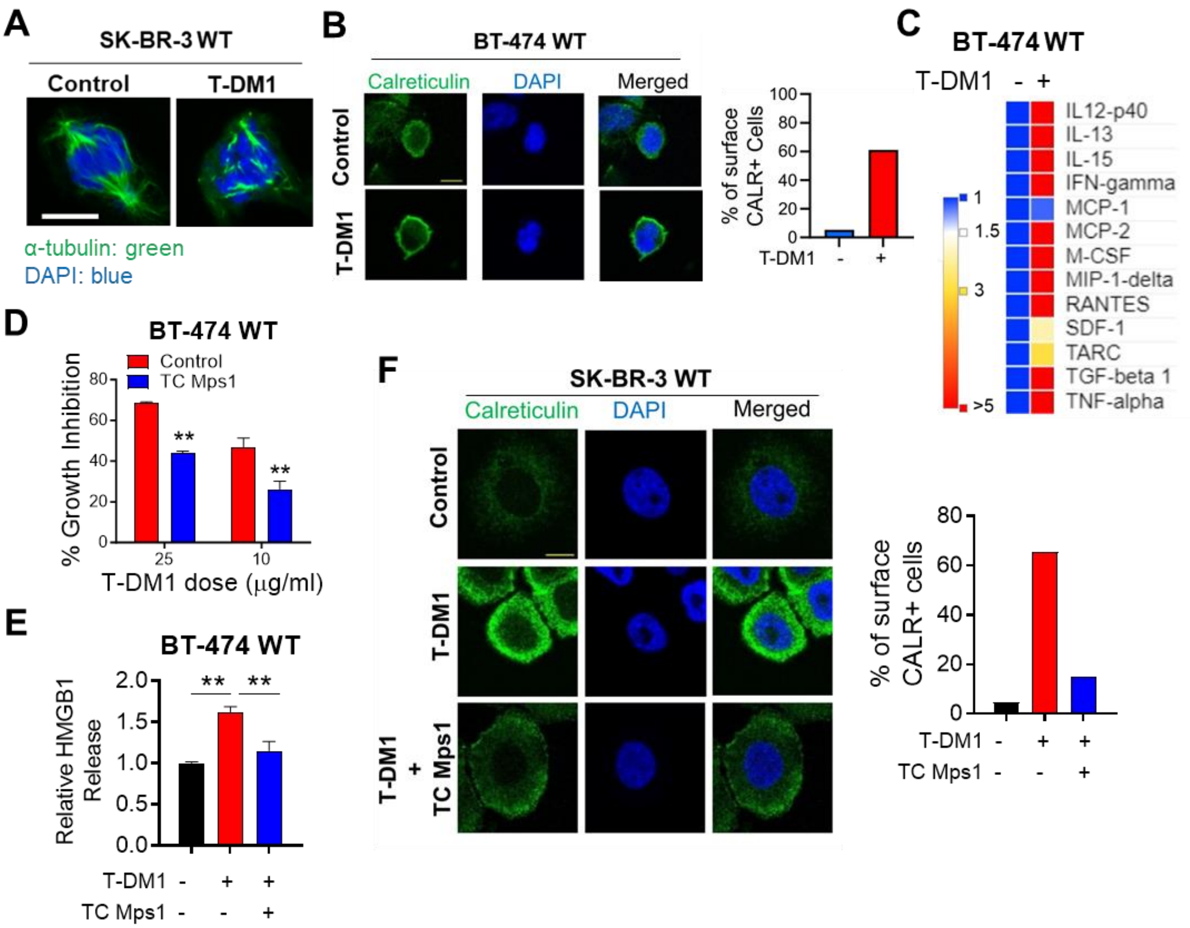

**Supplementary Figure S1. T-DM1 induces mitotic arrest and ICD markers in T-DM1 sensitive breast cancer cells in a SAC-dependent manner. A** IF staining of α-tubulin (green) in T-DM1 treated SK-BR-3 WT cell. Scale bar=10 µm. DAPI was used to stain the nucleus. **B** IF cell surface staining of calreticulin (green) in T-DM1 treated BT-474 WT cells. Scale bar=10 µm. Its quantification is provided on the right DAPI was used to stain the nucleus. **C** Chemokine array blot analysis inT-DM1 treated BT-474 WT cells. **D** Percent growth inhibition in BT-474 WT cells treated with T-DM1 alone or in combination with 1 µM TC Mps1 (Mps1 inhibitor) (n=3). **E** Relative HMGB1 release in BT-474 WT cells treated with T-DM1 alone or in combination with 1 µM TC Mps1 (n=3). **F** IF cell surface staining of calreticulin (green) in SK-BR-3 WT cells treated with T-DM1 alone or in combination with 1 µM TC Mps1. Scale bar=10 µm. Its quantification is provided on the right. Data correspond to mean values ± standard deviation (SD). *P*-values were calculated with the unpaired, two-tailed Student's t test. **, $P<0.01$.

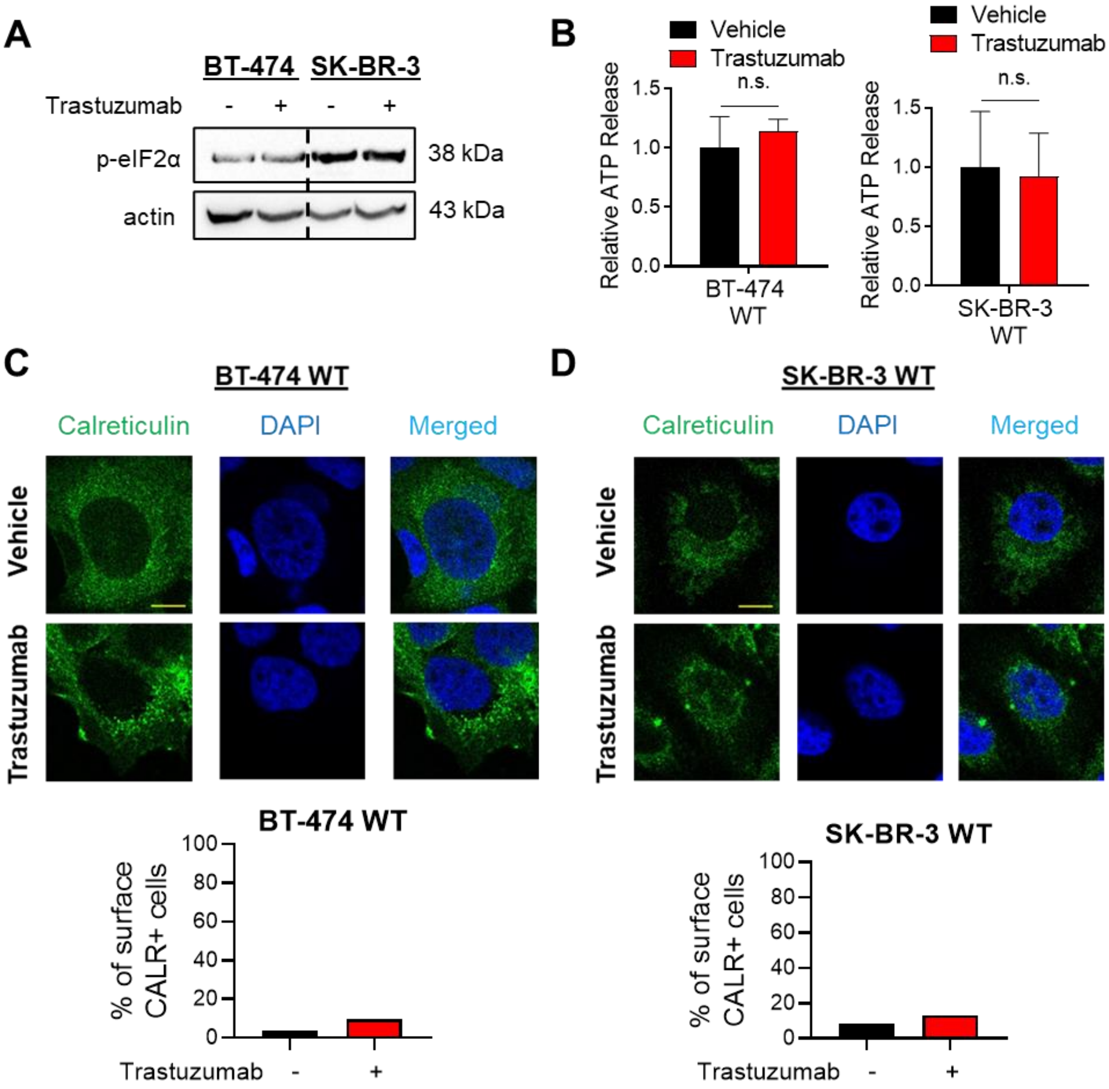

**Supplementary Figure S2. Effects of trastuzumab on ICD in BT-474 and SK-BR-3 WT cells. A** Western blot analysis of p-eIF2α in trastuzumab-treated SK-BR-3 and BT-474 WT cells. Actin is used as a loading control. **B** Relative ATP release in trastuzumab-treated SK-BR-3 WT and BT-474 WT cells (n=4, 5). **C, D** IF cell surface staining of calreticulin (green) in trastuzumab-treated BT-474 WT (C) and SK-BR-3 WT (D) cells. The quantification of percentage of surface CALR-positive cells are given below. Scale bar=10 µm. Data correspond to mean values ± standard deviation (SD). Significance was calculated with the unpaired, two-tailed Student's t test. n.s., not significant.

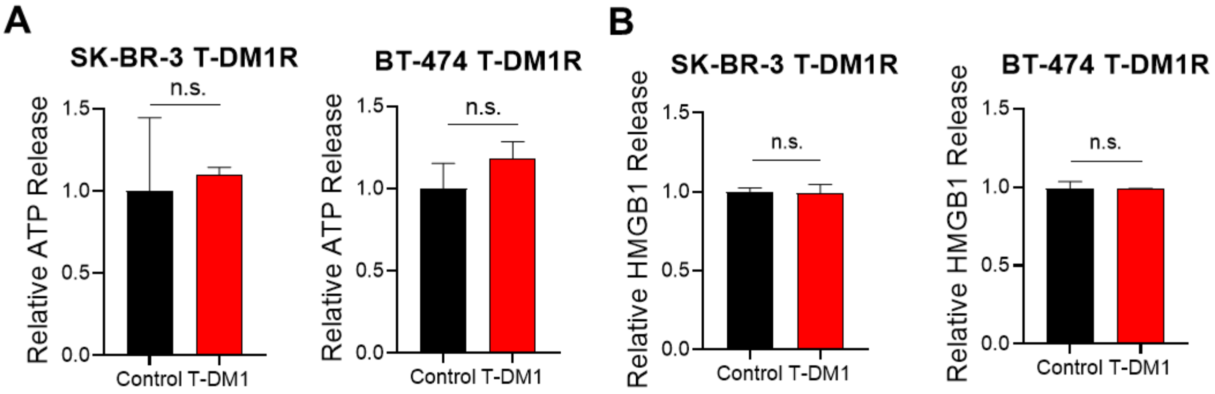

**Supplementary Figure S3. Effect of T-DM1 on ICD markers in T-DM1 resistant breast cancer cell lines. A, B** Relative ATP (A) and HMGB1 (B) release from T-DM1R cells treated with T-DM1 (n=3). Data correspond to mean values ± standard deviation (SD). Significance was calculated with the unpaired, two-tailed Student's t test. n.s., not significant.

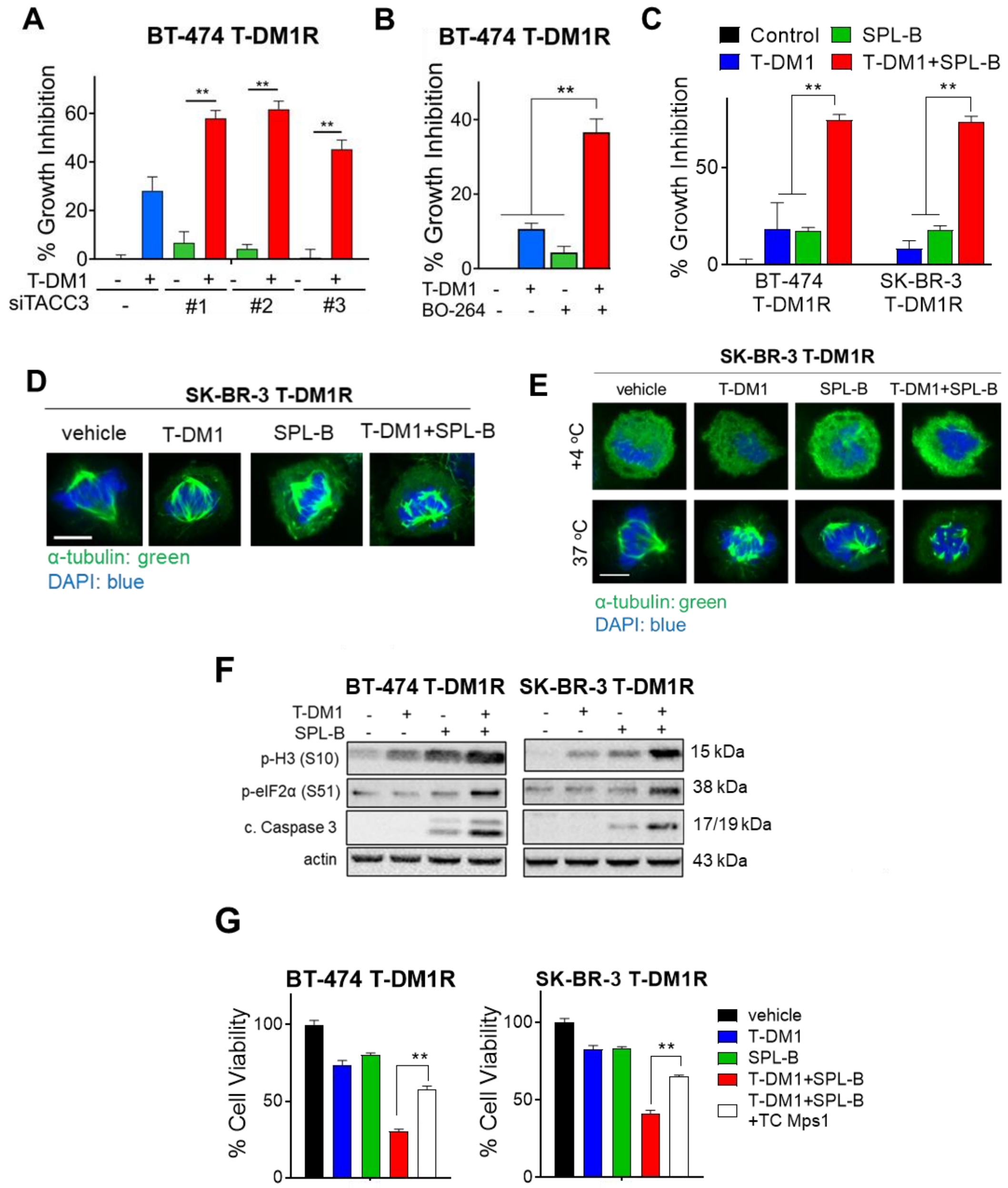

**Supplementary Figure S4. TACC3 inhibition overcomes T-DM1 resistance in a SAC-dependent manner. A** Percent growth inhibition in BT-474 T-DM1R cells transfected with siTACC3 and treated with 15 µg/mL T-DM1 (n=4-6). **B** Percent growth inhibition in BT-474 T-DM1R cells treated with T-DM1 alone or in combination with 1 µM TACC3 inhibitor (BO-264) (n=4-6). **C** Percentage growth inhibition in BT-474 and SK-BR-3 T-DM1R cells treated with T-DM1 alone or combination with 1.3 µM and 1.1 µM of the TACC3 inhibitor, SPL-B, respectively (n=3). **D** IF staining of α-tubulin (green) in SK-BR-3 T-DM1R cells treated with T-DM1 or SPL-B alone or combination. Scale bar=10 µm. **E** Microtubule polymerization assay in SK-BR-3 T-DM1R cells treated with T-DM1 or SPL-B alone or combination. Scale bar=10 µm. **F** Western blot analysis of mitotic arrest, apoptosis and ICD markers in T-DM1R cells treated with T-DM1 alone or in combination with SPL-B. Actin is used as a loading control. **G** Percentage cell viability in BT-474 and SK-BR-3 T-DM1R cells treated with combination therapy with or without TC Mps1 (n=4, 5). Data correspond to mean values ± standard deviation (SD). *P*-values were calculated with the unpaired, two-tailed Student's t test. **, *P*<0.01.

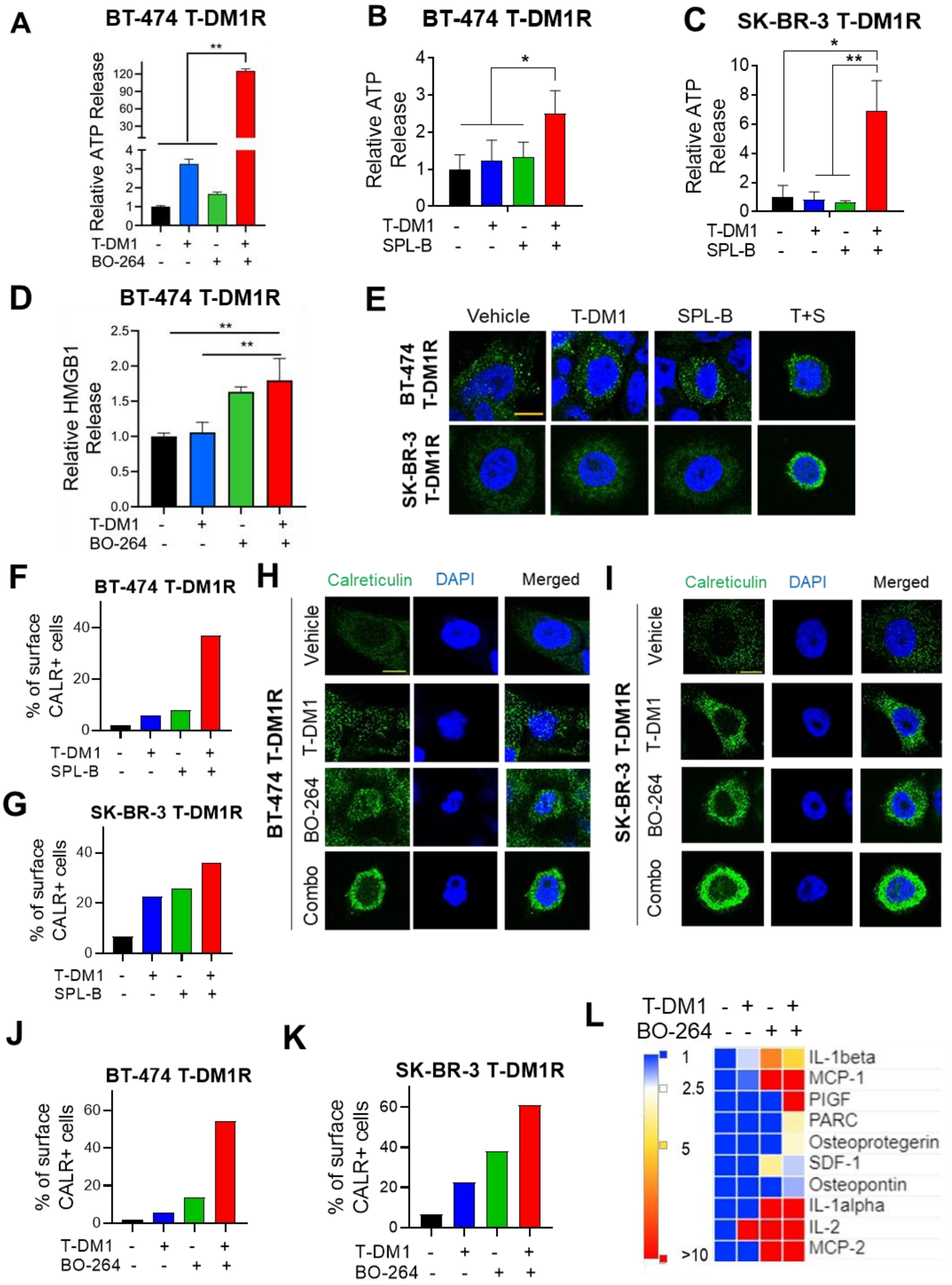
A
BT-474 T-DM1R
Relative ATP Release
T-DM1
BO-264
B
BT-474 T-DM1R
Relative ATP Release
T-DM1
SPL-B
C
SK-BR-3 T-DM1R
Relative ATP Release
T-DM1
SPL-B
D
BT-474 T-DM1R
Relative HMGB1 Release
T-DM1
BO-264
E
Vehicle
T-DM1
SPL-B
T+S
BT-474 T-DM1R
SK-BR-3 T-DM1R
F
BT-474 T-DM1R
% of surface CALR+ cells
T-DM1
SPL-B
G
SK-BR-3 T-DM1R
% of surface CALR+ cells
T-DM1
SPL-B
H
Calreticulin
DAPI
Merged
BT-474 T-DM1R
Vehicle
T-DM1
BO-264
Combo
I
Calreticulin
DAPI
Merged
SK-BR-3 T-DM1R
Vehicle
T-DM1
BO-264
Combo
J
BT-474 T-DM1R
% of surface CALR+ cells
T-DM1
BO-264
K
SK-BR-3 T-DM1R
% of surface CALR+ cells
T-DM1
BO-264
L
T-DM1
BO-264
1
2.5
5
>10
IL-1beta
MCP-1
PIGF
PARC
Osteoprotegerin
SDF-1
Osteopontin
IL-1alpha
IL-2
MCP-2

**Supplementary Figure S5. Inhibition of TACC3 in combination with T-DM1 restores ICD markers in T-DM1R cells in vitro. A** Relative ATP release from BT-474 T-DM1R cells treated with T-DM1 alone or in combination with BO-264 (n=4). **B, C** Relative ATP release from BT-474 (B) and SK-BR-3 (C) T-DM1R cells treated with T-DM1 alone or in combination with SPL-B (n=3, 4). **D** Relative HMGB1 release from BT-474 T-DM1R cells treated with T-DM1 alone or in combination with BO-264 (n=3). **E** IF cell surface staining of calreticulin (green) in T-DM1R cells treated with T-DM1 alone or in combination with SPL-B. Scale bar=10 µm. **F, G** The quantification of percentage of surface CALR-positive cells from E. **H, I** IF cell surface staining of calreticulin (green) in T-DM1R cells treated with T-DM1 alone or in combination with BO-264. Scale bar=10 µm. **J, K** The quantification of percentage of surface CALR-positive cells from H, I. **L.** Cytokine array analysis showing relative changes in secreted cytokines among single agent- and combination-treated SK-BR-3 T-DM1R cells. Data correspond to mean values ± standard deviation (SD). *P*-values were calculated with the unpaired, two-tailed Student's t test. *, $P<0.05$; **, $P<0.01$.

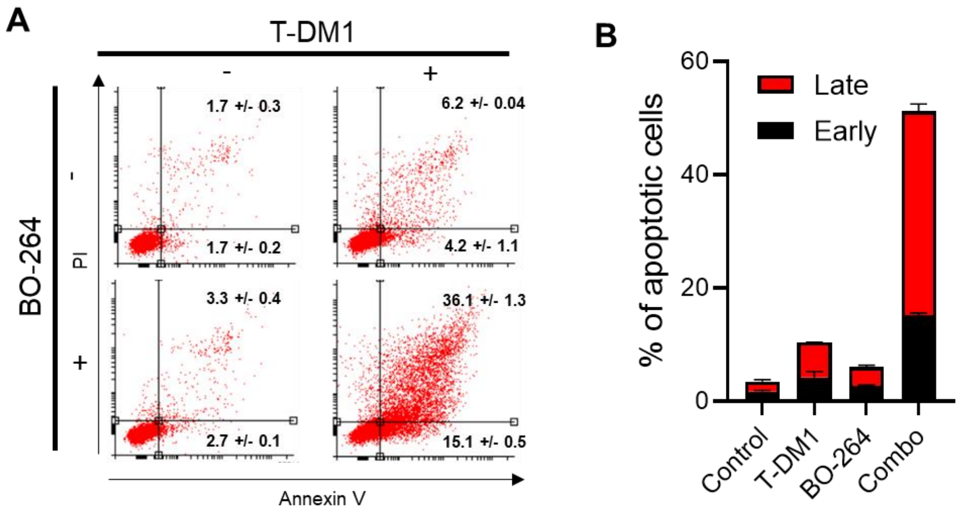

**Supplementary Figure S6. T-DM1 combination with BO-264 induces apoptosis in T-DM1 resistant EMT6.huHER2 murine mammary tumor cell line with human HER2. A, B** Annexin V/PI staining of EMT6.huHER2 cells treated with T-DM1 or BO-264 or their combination (A) and its quantification (B) (n=2).

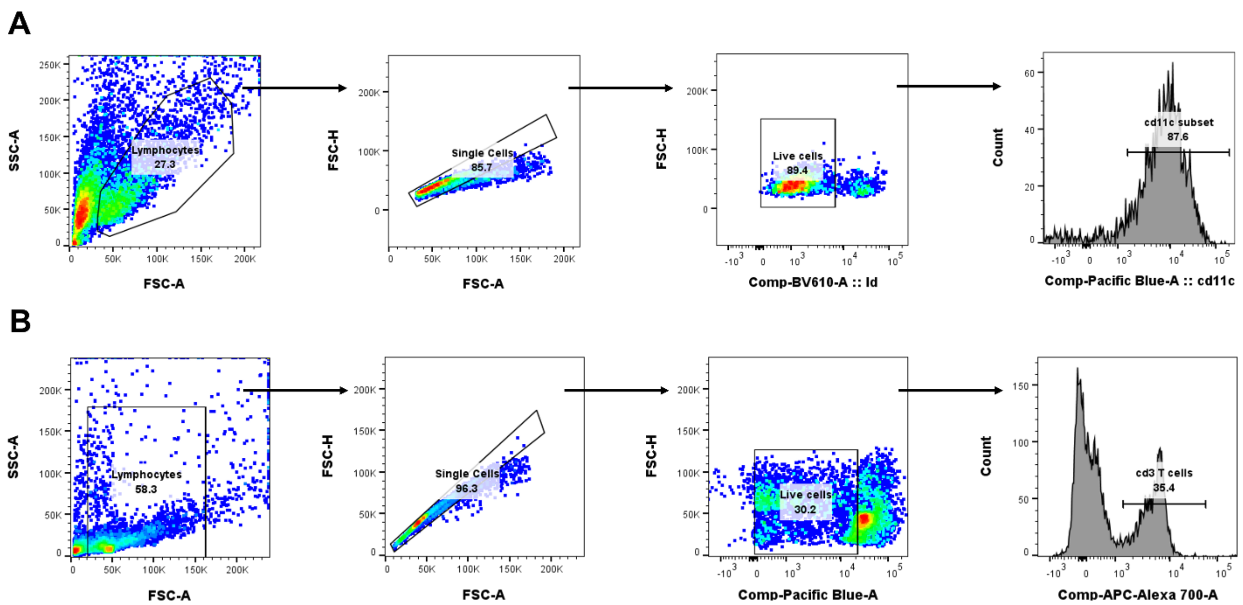

**Supplementary Figure S7. Gating strategy for flow analysis A, B** Gating strategy for CD80/CD86 (A) and CD8/CD25 double positive cells (B) for flow cytometry analysis.

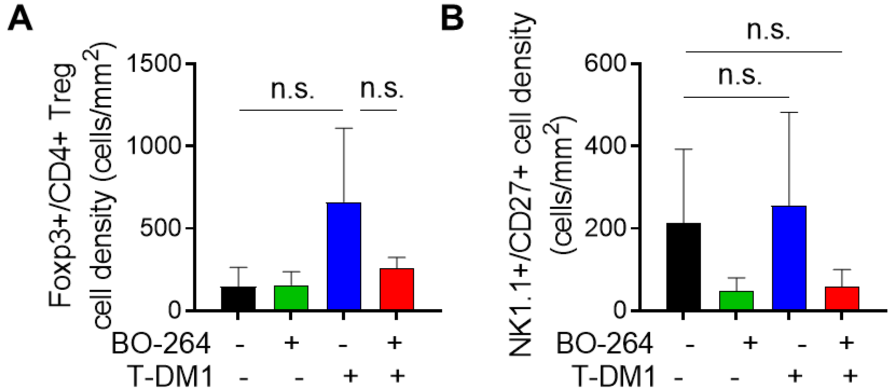

**Supplementary Figure S8. Infiltration of Treg and NK cells into combination treated MMTV.f.huHER2#5 tumors. A, B** Quantification of Treg (A) and NK (B) cell densities within MMTV.f.huHER2#5 tumors from mice treated with T-DM1 alone or in combination with BO-264 (n=3). Data correspond to mean values ± standard deviation (SD). Significance was calculated with the unpaired, two-tailed Student's t test. n.s., not significant.

### Supplementary Tables

**Supplementary Table S1.** Sequences of the siRNAs used.

| Gene Name | NCBI Gene ID | Company | Catalog number | sequence |
|---|---|---|---|---|
| TACC3 | 10460 | Dharmacon | D-004155-02-0005 | GAGCGGACCUGUAAAACUA |
| TACC3 | 10460 | Dharmacon | D-004155-03-0005 | GAACGAAGAGUCACUGAAG |
| TACC3 | 10460 | Dharmacon | D-004155-04-0005 | UCUCUUAGGUGUCAUGUUC |

**Supplementary Table S2.** List of antibodies used in Western blot (WB), immunofluorescence (IF) and flow cytometry (F) experiments.

| Antibody | Provider | Catalog number | WB dilution | IF dilution | F dilution |
|---|---|---|---|---|---|
| Beta-actin | Santa Cruz | sc-8432 | 1:10000 | - | - |
| TACC3 | Santa Cruz | sc-376883 | 1:1000 | - | - |
| PARP | Cell Signaling | 9542S | 1:1000 | - | - |
| eIF2α | Santa Cruz | sc-133132 | 1:1000 | - | - |
| Phospho-eIF2α (Ser51) | Cell Signaling | 9721 | 1:1000 | - | - |
| Phospho-Histone H3 (Ser10) | Cell Signaling | 9701 | 1:1000 | - | - |
| Cleaved caspase 3 | Cell Signaling | 9664 | 1:1000 | - | - |
| Cleaved PARP | Cell Signaling | 5625 | 1:1000 | - | - |
| PARP | Cell Signaling | 9542 | 1:1000 | - | |
| Cyclin B1 | Cell Signaling | 4138 | 1:1000 | - | - |
| CD11c | Cell Signaling | 97585 | - | 1:50 | - |
| CD86 | Cell Signaling | 19589 | - | 1:50 | - |
| CD27 | Abcam | ab175403 | - | 1:100 | - |
| NK 1.1 | Cell Signaling | 39197 | - | 1:50 | - |
| CD4 | Abcam | ab183685 | - | 1:100 | - |
| CD8 | Cell Signaling | 98941 | - | 1:50 | - |
| Foxp3 | Cell Signaling | 12653 | - | 1:50 | - |
| CD25 | Cell Signaling | 39475 | - | 1:50 | - |
| CD3 | Cell Signaling | 85061 | - | 1:50 | - |
| Calreticulin (Calregulin) | Santa Cruz | sc-373863 | - | 1:500 | - |
| Alpha-tubulin | Santa Cruz | sc-32293 | - | 1:1000 | - |
| Anti-mouse IgG | Cell Signaling | 7076 | 1:3000 | - | - |
| Anti-rabbit IgG | Cell Signaling | 7074 | 1:3000 | - | - |
| Alexa Fluor® 488 anti-mouse | Life Technologies | A-11001 | - | 1:1000 | - |
| Alexa Fluor® 647 anti-rabbit | Life Technologies | A-31573 | - | 1:1000 | - |
| DAPI | Life Technologies | D1306 | - | 1:5000 | - |

| | | | | | |
|---|---|---|---|---|---|
| CD3 (Alexa Alexa Fluor® 488) | Biolegend | 100321 | - | - | 1:500 |
| CD8 (PE/Cy7) | Biolegend | 100722 | - | - | 1:500 |
| CD80 (APC) | eBioscience | 17-0801-82 | - | - | 1:500 |
| CD86 (PE/Cy7) | Biolegend | 105103 | - | - | 1:500 |
| CD25 (Brilliant Violet 510) | Biolegend | 102042 | - | - | 1:500 |
